## supplemental file for "Detecting Regime Shifts: Neurocomputational Substrates for Over- and Underreactions to Change"

#### Supplementary Methods

##### Experiment 2

The procedure of Experiment 2 was identical to Experiment 1, except that there was no regime shift in this experiment. In other words, we only manipulated the signal diagnosticity in this experiment. The regime that the sensory signals were drawn from (red or blue urn) was randomly determined such that in half of the trials the regime was red and the other half the regime was blue. The order of the regimes across trials was pseudo randomized for each subject separately. Identical to Experiment 1, the subjects in Experiment 2 were instructed to estimate the probability that regime was the blue urn at the presentation of each new signal. Therefore, in both Experiments 1 and 2, the subjects estimated the probability that the current regime was the blue regime. However, it was only in Experiment 1 that the probability estimates conveyed information about the subjects' belief about change (whether the regime had shifted).

##### Experiment 3

We referred to Experiment 3 as the motor-equivalent task of Experiments 1 and 2. The primary goal of this experiment was to rule out the motor confounds for probability estimates in Experiment 1. We were aware that brain regions whose activity correlated with probability estimates in Experiment 1 can simply reflect entering of numbers through button presses and thus have nothing to do with estimating the probability of change. Therefore, to establish the evidence for the neural representations of probability estimates, we needed to rule out the motor confounds. The motor-equivalent task shared many of the physical and motor aspects of the tasks in Experiments 1 and 2. First, the task structure was identical. In the motor-equivalent task, each trial contained ten rounds. In each round, a hollow dot with no color information was shown to indicate the number of this round. A random number from 0 to 99 was presented at the center of screen. Subjects had to enter the indicated number with two successive button presses within 4 seconds. After this button-press stage there was feedback on the number they just entered, with white given for correct inputs and yellow given for incorrect entries. The reward was $3 for each correct answer and -$3 for incorrect answer. The accumulated reward outcome for each trial was revealed at the end of the trial.

#### Computational models for Experiment 2

Below we describe the computational models (Bayesian model and system-neglect model) for Experiment 2 where subjects performed identical task as in Experiment 1 with one exception that no regime shift was possible (transition probability $q = 0$). We fit the system-neglect model and the parameter estimates can be found in Supplementary Fig. 2.

##### Bayesian model

The Bayesian posterior odds of the blue regime at the *t*-th period given the signal history $H_t$ were calculated by:

$$\frac{P_t^B}{1-P_t^B} = \frac{\Pr(B_t|H_t)}{\Pr(R|H_t)} = \frac{\Pr(B)}{\Pr(R)} \times \frac{\Pr(H_t|B)}{\Pr(H_t|R)}.$$

where $\frac{\Pr(B)}{\Pr(R)}$ is the prior odds and $\frac{\Pr(H_t|B)}{\Pr(H_t|R)}$ is the likelihood ratio. Given that the probability of being in the blue regime and being in the red regime were equal $\Pr(B) = \Pr(R) = 0.5$, Eq. (3) can be rewritten as

$$\frac{P_t^B}{1-P_t^B} = \frac{\Pr(B_t|H_t)}{\Pr(R|H_t)} = \frac{0.5}{0.5} \times \frac{\Pr(H_t|B)}{\Pr(H_t|R)} = \frac{\Pr(H_t|B)}{\Pr(H_t|R)}.$$

The likelihood ratio is computed according to the following equation

$$\frac{\Pr(H_t|B)}{\Pr(H_t|R)} = d^{t-2\sum_{k=1}^{t} r_k}.$$

where *d* represents signal diagnosticity. The exponent of *d* captures the difference between the number of the blue balls and red balls from the first to the *t*-th period. For example, suppose that at the 5th period there are 4 red balls and 1 blue ball. Then $d^{t-2\sum_{k=1}^{t} r_k}$ would be $d^{5-2\times 3} = d^{-3}$ where the exponent is the number of blue balls minus the number of red balls.

##### System-neglect model

Similar to Experiment 1, we developed and fit the system-neglect model. The system-neglect model consisted of a weighting parameter $\beta$ for signal diagnosticity

$$\frac{P_t^B}{1-P_t^B} = \frac{\Pr(B_t|H_t)}{\Pr(R|H_t)} = d^{\beta(t-2\sum_{k=1}^{t} r_k)}.$$

where we separately estimated $\beta$ for each level of signal diagnosticity

$$\beta = \beta_1 D_1 + \beta_2 D_2 + \beta_3 D_3$$

Here $D_n$ is the dummy variable for diagnosticity for $d_n$. Note that $\beta = 1$ corresponds to Bayesian estimates.

#### SUPPLEMENTARY RESULTS

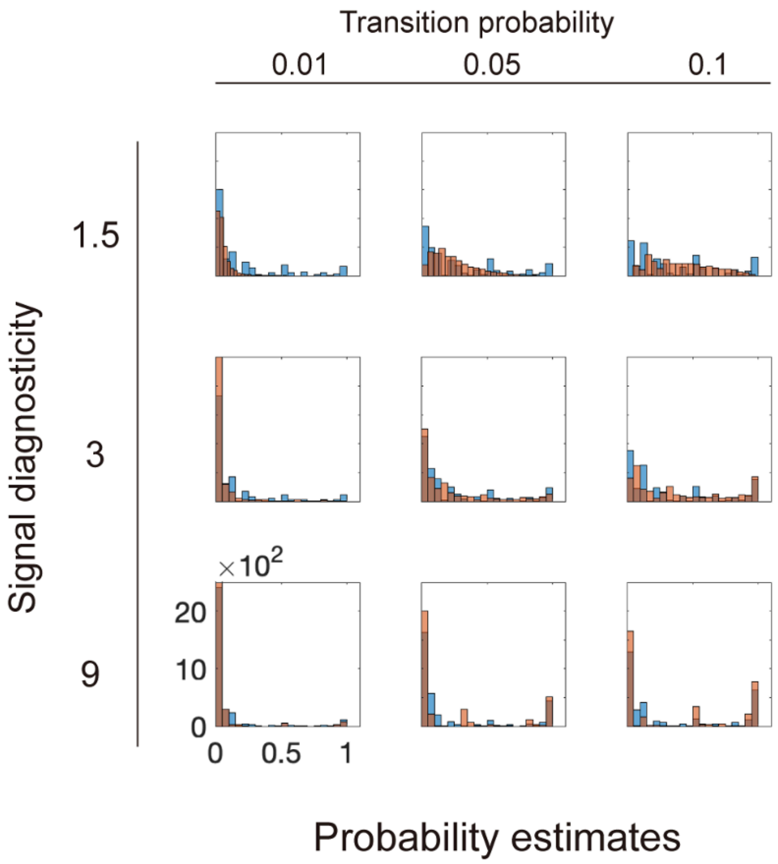


**Figure S1.** Probability estimates from all subjects are plotted as histograms separately for each condition—a combination of transition probability and signal diagnosticity. The blue bars represent the actual probability estimates, while the orange bars correspond to the probability estimates predicted by the Bayesian model.

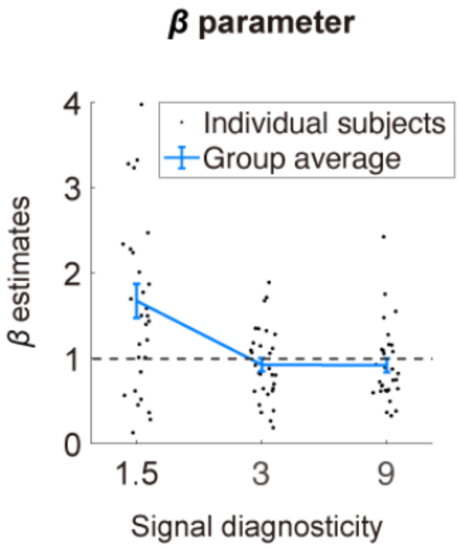


**Figure S2.** Experiment 2: estimates of the weighting parameter for signal diagnosticity $(\beta)$ in the system-neglect model. Dashed lines indicate parameter value equal to 1.

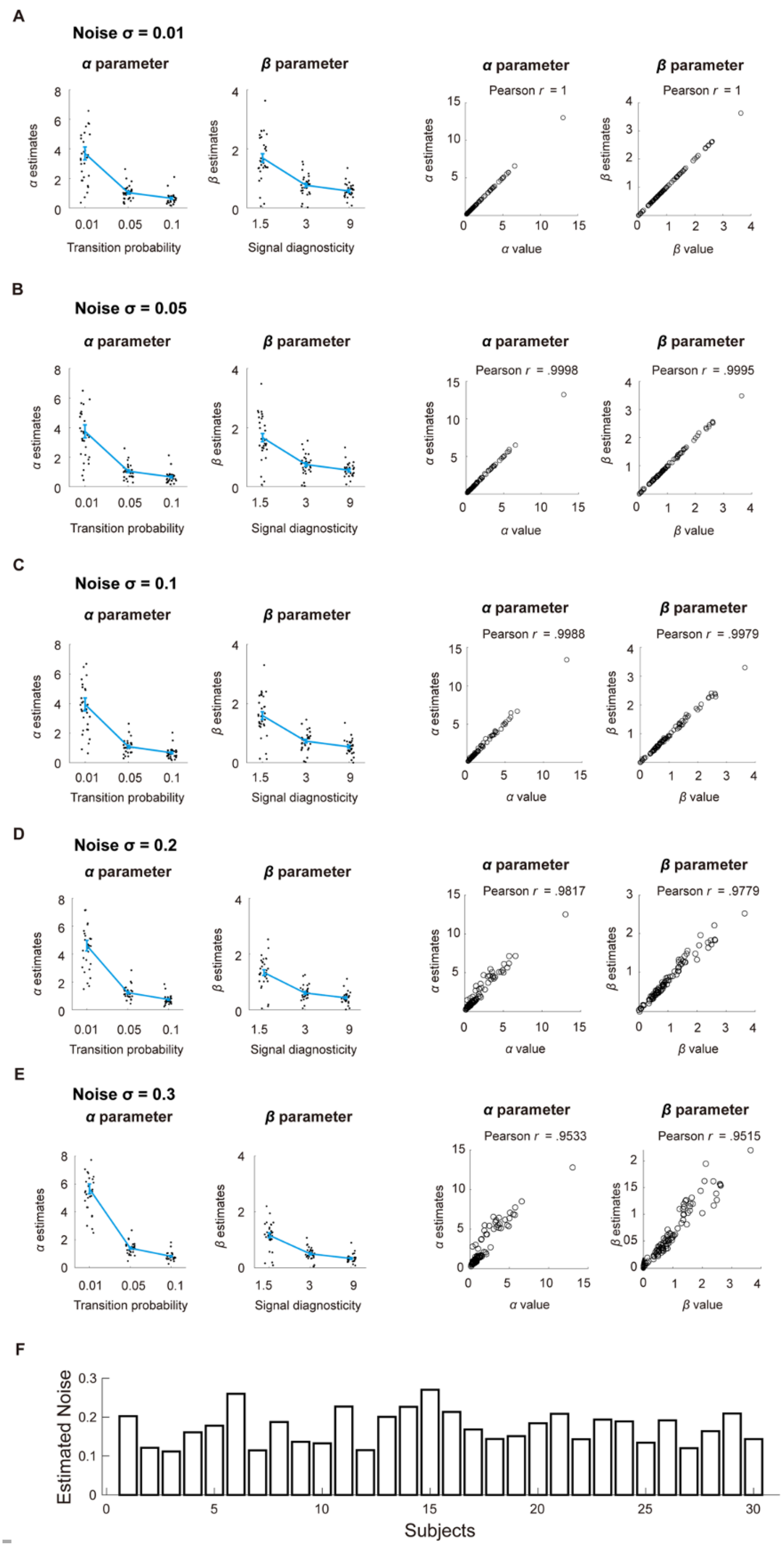
A
Noise σ = 0.01
α parameter
β parameter
α estimates
β estimates
Transition probability
Signal diagnosticity
Pearson r = 1
α value
β value
B
Noise σ = 0.05
Pearson r = .9998
Pearson r = .9995
C
Noise σ = 0.1
Pearson r = .9988
Pearson r = .9979
D
Noise σ = 0.2
Pearson r = .9817
Pearson r = .9779
E
Noise σ = 0.3
Pearson r = .9533
Pearson r = .9515
F
Estimated Noise
Subjects

**Figure S3.** Parameter recovery. We simulated probability estimates according to the system-neglect model. We used each subject's parameter estimates as our choice of parameter values used in the simulation. Using simulated data, we estimated the parameters ($\alpha$ and $\beta$) in the system-neglect model. To examine parameter recovery, we plotted the parameter values we used to simulate the data against the parameter estimates we obtained based on simulated data and computed their Pearson correlation. Further, we added different levels of Gaussian white noise with standard deviation $\sigma = [0.01, 0.05, 0.1, 0.2, 0.3]$ to the simulated data to examine parameter recovery and show the results respectively in Fig. A, B, and C. For each noise level, we show the parameter estimates in the left two graphs. In the right two graphs, we plot the parameter estimates based on simulated data against the parameter values used to simulate the data. **A.** Noise $\sigma = 0.01$. **B.** Noise $\sigma = 0.05$. **C.** Noise $\sigma = 0.1$. **D.** Noise $\sigma = 0.2$. **E.** Noise $\sigma = 0.3$. **F.** Empirically estimated noise ($\sigma$) of each subject. Each bar represents a subject's estimated noise level.

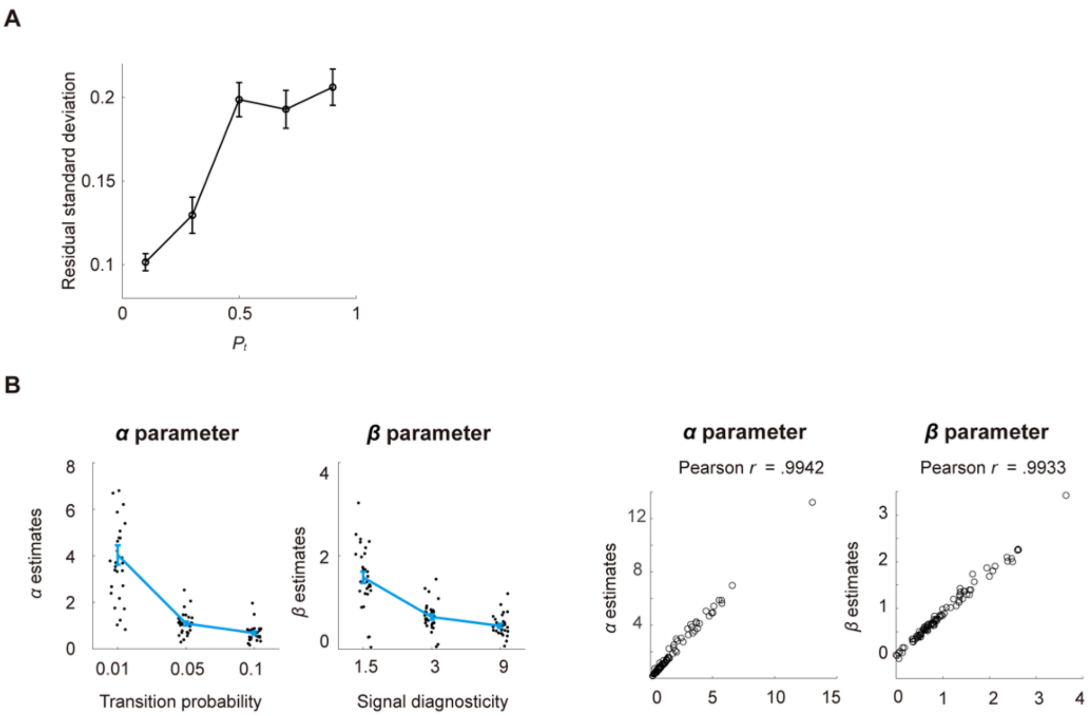


**Figure S4.** Impact of noise homoscedasticity on parameter estimation. **A.** Empirically estimated residual standard deviation. Mean residual standard deviation (across subjects, black data points) in the five probability intervals, [0.0–0.2), [0.2–0.4), [0.4–0.6), [0.6–0.8), and [0.8–1.0], were 0.1015, 0.1296, 0.1987, 0.1929, and 0.2061, respectively. Error bars represent $\pm 1$ standard error of the mean. **B.** Parameter recovery results assuming heteroscedastic noise. We performed parameter recovery using the empirically estimated, probability-dependent residual variance shown in A (the mean residual standard deviation estimates). Conventions are the same as in Fig. S3.

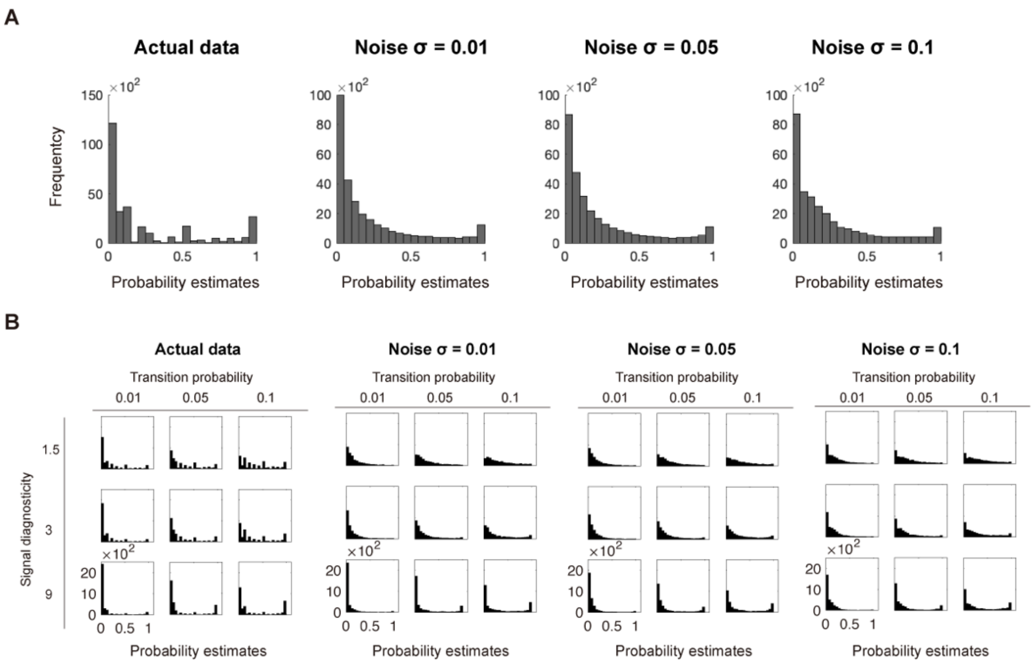


**Figure S5.** Probability estimates from the actual and simulated data. **A.** Histogram of subjects' actual probability estimation data collapsed across all conditions (left graph) and simulated probability estimation data under three different noise levels (Noise $\sigma = 0.01, 0.05, 0.1$). Descriptions of how we performed simulations can be seen in Supplementary Fig. S3. **B.** Subjects data are plotted as histograms separately for each condition.

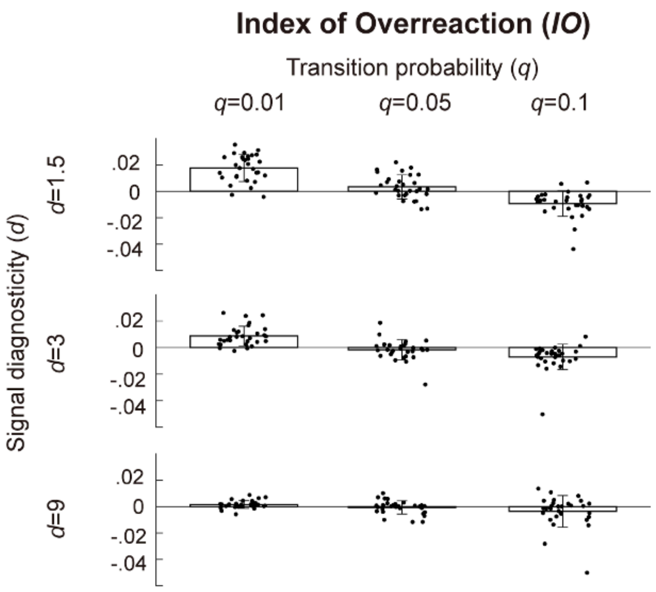


**Figure S6.** System-neglect model can well-describe subjects' over- and underreactions to change. We fit the system-neglect model to each individual subject' probability estimates and used the resulting parameter estimates to compute each subject's probability estimates under the system-neglect model ($P_t^{SN}$). We then used $P_t^{SN}$ to compute Index of Overreaction ($IO$). Here, $IO$ was computed by subtracting belief revision predicted by the Bayesian model ($\Delta P_t^B = P_t^B - P_{t-1}^B$) from belief revision estimated by system-neglect model ($\Delta P_t^{SN} = P_t^{SN} - P_{t-1}^{SN}$). Formally, $IO = \Delta P_t^{SN} - \Delta P_t^B$. The mean $IO$ (across all subjects; indicated by the bars) is plotted as a function of transition probability and signal diagnosticity. Solid symbols represent data points from individual subjects. Error bars represent $\pm 1$ standard error of the mean. The patterns of over- and underreactions here resembled those based on actual data (Fig. 2B), suggesting that the system-neglect model can describe subjects' over- and underreactions well.

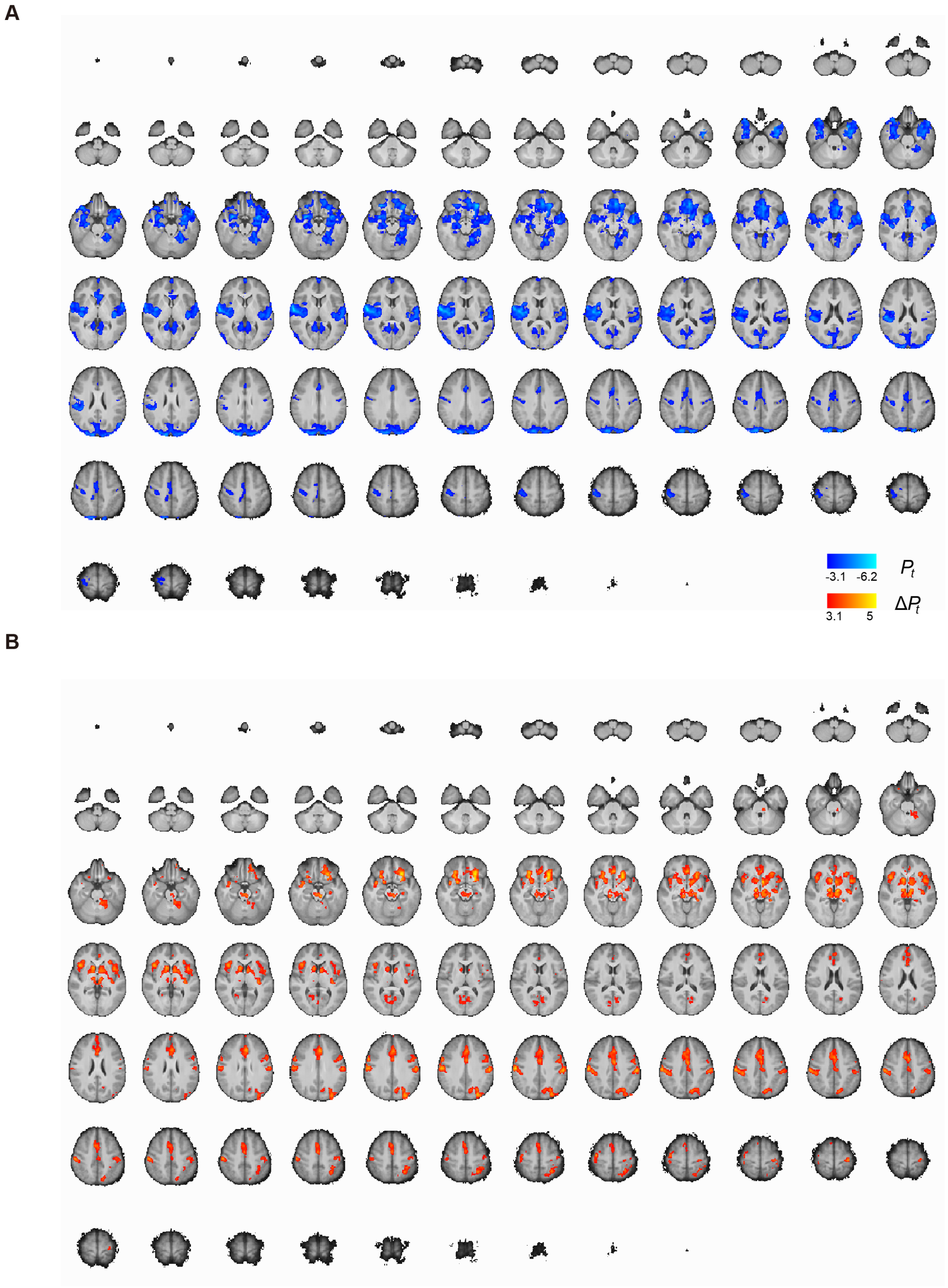


**Figure S7.** Whole-brain results of GLM-1 on the main experiment (Experiment 1) showing brain regions that significantly correlate with regime-shift probability estimates ($P_t$; clusters in blue in Panel A) and the updating of beliefs about change ($\Delta P_t$; clusters in orange in Panel B). Cluster-level inference using Gaussian random field theory (familywise error-corrected at $p < .05$ using with a cluster-forming threshold $z > 3.1$).

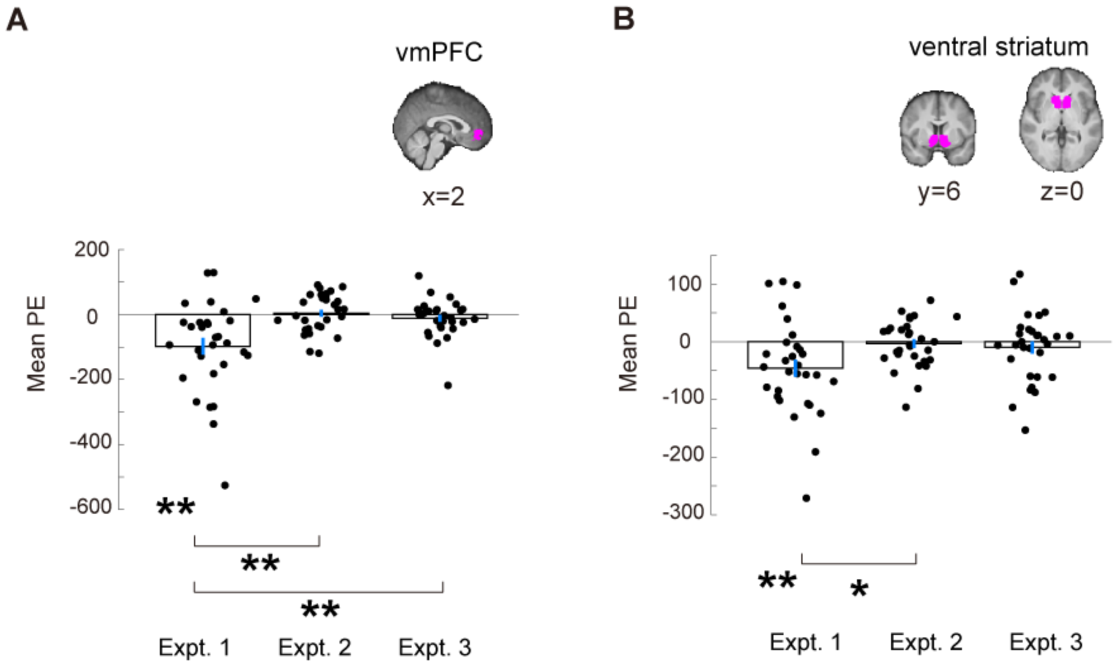


**Figure S8.** Independent region-of-interest (ROI) analysis in vmPFC and ventral striatum on probability estimates across the three experiments (Experiment 1 was the main experiment on regime-shift detection, Experiments 2 and 3 were control experiments). The vmPFC ROI we used was based on Bartrat et al. (2013). The ventral striatum mask was based on nucleus accumbens mask from Harvard-Oxford Cortical Structural Atlas. For each subject and each ROI, we extracted the mean parameter estimates (PE) of the probability estimates contrast in GLM-1. **A.** vmPFC ROI. Experiment 1: One-sample *t* test, $t(29) = -3.82,\ p < 0.01$; Experiment 2: One-sample *t* test, $t(29) = 0.36,\ p = 0.71$; Experiment 3: One-sample *t* test, $t(29) = -1.11,\ p = 0.28$; Experiments 1 $-$ Experiment 2: two-sample *t* test, $t(58) = -3.67, p < 0.01$; Experiments 1 $-$ Experiment 3: two-sample *t* test, $t(58) = -3.12,\ p < 0.01$;. **B.** Ventral striatum ROI. Experiment 1: $(29) = -3.06,\ p < 0.01$; Experiment 2: $t(29) = 0.44,\ p = 0.67$; Experiment 3: $t(29) = -0.93,\ p = 0.36$; Experiments 1 $-$ Experiment 2: $t(58) = -2.55,\ p = 0.01$; Experiments 1 $-$ Experiment 3: $t(58) = -1.95,\ p = 0.06$. The * symbol indicates $p$<0.05 (two-tailed), and ** symbol indicates $p$<0.01 (two-tailed).

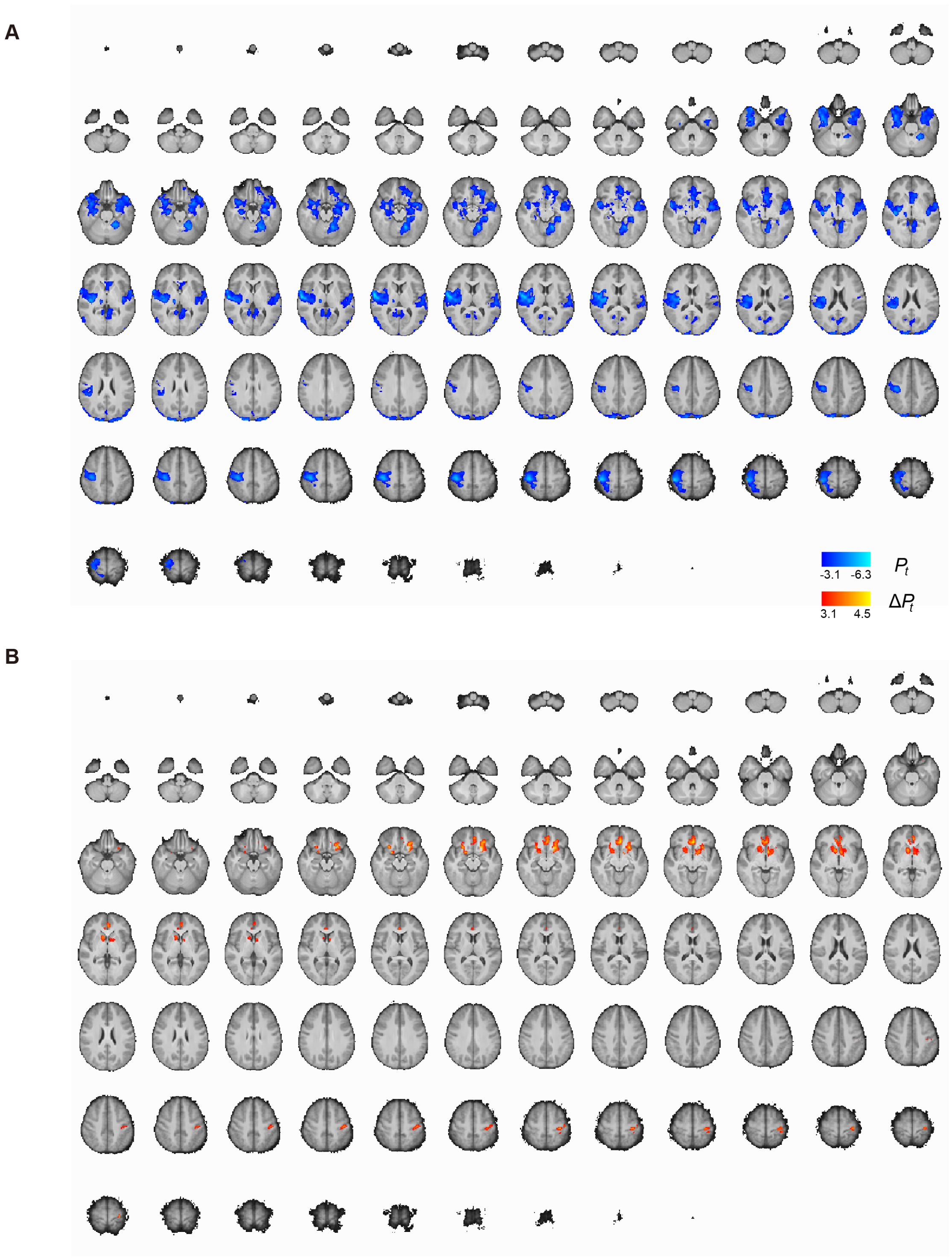


**Figure S9.** Whole-brain results of GLM-2 on the main experiment (Experiment 1) showing brain regions that significantly correlate with regime-shift probability estimates ($P_t$; clusters in blue in Panel A) and the updating of beliefs about change ($\Delta P_t$; clusters in orange in Panel B). Cluster-level inference using Gaussian random field theory (familywise error-corrected at $p < .05$ using with a cluster-forming threshold $z > 3.1$).

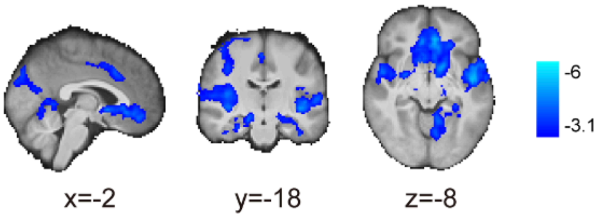


**Figure S10.** Whole-brain results on activity that significantly correlated with the subjects' log odds estimates of regime shift, $\ln(P_t/(1-P_t))$, where $P_t$ represents subjects' probability estimates. In this analysis, we replaced the parametric regressor of $P_t$ with the log odds of regime shifts in GLM-1. Color for significant activations: blue indicates negative correlation with $\ln(P_t/(1-P_t))$. Familywise error-corrected at $p < .05$ using Gaussian random field theory with a cluster-forming threshold $z > 3.1$.

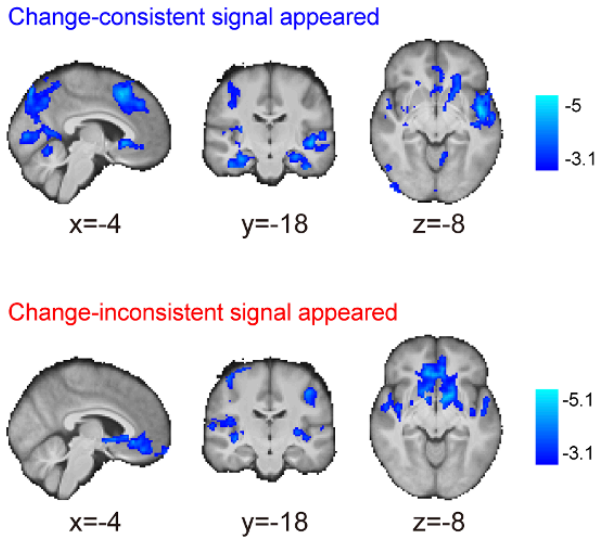


**Figure S11.** Probability estimates of regime shifts ($P_t$) separately analyzed for change-consistent (blue) and change-inconsistent (red) signals. Whole-brain results on activity that significantly correlated with probability estimates of regime shift for change-consistent and change-inconsistent signals. We observed that vmPFC correlated with $P_t$ for both types of signals. Blue color showing significant activations indicates negative correlation with $P_t$. Familywise error-corrected at $p < .05$ using Gaussian random field theory with a cluster-forming threshold $z > 3.1$.

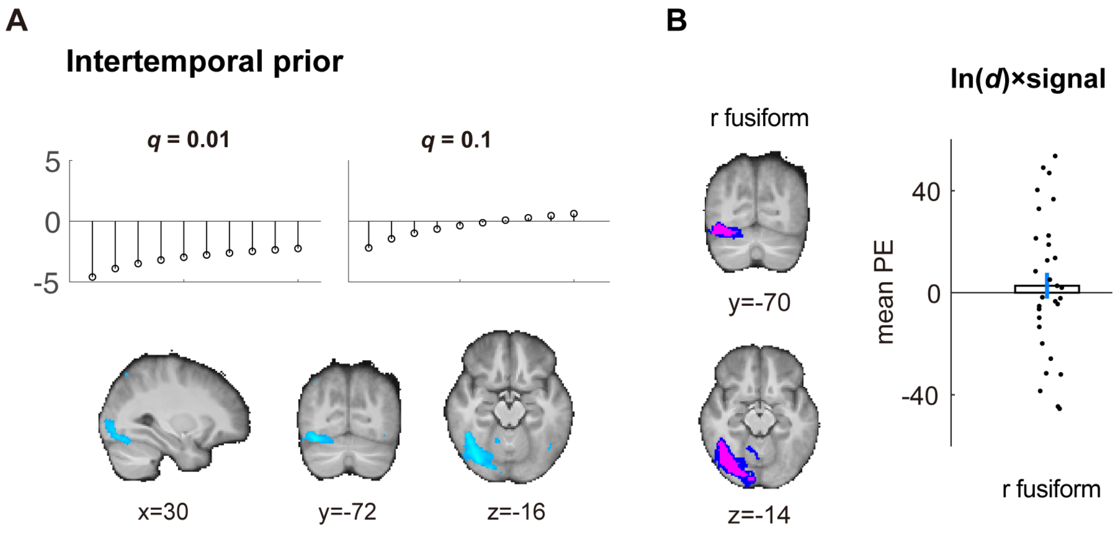


**Figure S12.** Key variables for regime-shift computations. **A.** Whole-brain results on the intertemporal prior of regime shift. **B.** Using the intertemporal prior ROI (left graph: magenta indicates voxels shared by the LOSO ROI of all subjects; blue indicates voxels of LOSO ROI of at least one subject) to examine the regression coefficients of the strength of evidence in favor of change, $\ln(d) \times$ signal. The mean parameter estimates (mean PE), i.e., regression coefficient, was not significantly different from 0 (one-sample $t$ test, $t(29) = 0.54, p = 0.59$, two-tailed).

**Table S1.** Experiment 1: Probability estimates $(P_t)$ and belief revision $(\Delta P_t)$ contrasts based on GLM-1. Cluster-level inference using Gaussian random field theory (familywise error corrected at $p < .05$ with a cluster-forming threshold $z > 3.1$).

| **Probability estimates $P_t$ (negative correlation)** | | | | |
|---|---|---|---|---|
| **Cluster** | **Hemisphere** | **Cluster size** | **z-max** | **z-max(x,y,z)** |
| Central Opercular Cortex | R | 26990 | 6.13 | (62,-6,6) |
| (Local maxima) | | | | |
| Frontal Orbital Cortex | | | 5.80 | (−28, 32, −14) |
| Insular Cortex | | | 5.72 | (36, −14, 16) |
| Planum Polare | | | 5.58 | (−48, −8, −10) |
| Central Opercular Cortex | | | 5.53 | (50, −10, 12) |
| Frontal Orbital Cortex | | | 5.45 | (30, 32, -16) |
| Cingulate Gyrus, anterior division | - | 989 | 4.51 | (0,18,36) |
| (Local maxima) | | | | |
| Cingulate Gyrus, posterior division | | | 3.94 | (14, −34, 46) |
| Cingulate Gyrus, anterior division | | | 3.86 | (−8, −10, 40) |
| Cingulate Gyrus, anterior division | | | 3.86 | (6, 0, 40) |
| Supplementary Motor Cortex | | | 3.84 | (6, −8, 48) |
| Supplementary Motor Cortex | | | 3.81 | (6, −4, 48) |
| **Belief revision $\Delta P_t$ (positive correlation)** | | | | |
| Frontal Orbital Cortex | L | 3105 | 4.94 | (-24,18,-10) |
| Cingulate Gyrus, anterior division | L | 2589 | 4.34 | (-4,26,30) |
| Frontal Orbital Cortex | R | 1796 | 4.57 | (26,20,-12) |
| Right Cerebral White Matter | R | 1557 | 4.67 | (12,-14,-4) |
| Lateral Occipital Cortex, superior division | L | 1366 | 4.34 | (-28,-82,34) |

| | | | | |
|---|---|---|---|---|
| Postcentral Gyrus | R | 1262 | 4.71 | (60,-18,36) |
| Postcentral Gyrus | L | 1088 | 4.48 | (-58,-22,36) |
| Postcentral Gyrus | L | 587 | 4.25 | (-34,-26,68) |

**Table S2.** Experiment 1: Probability estimates $(P_t)$ and belief revision $(\Delta P_t)$ contrasts based on GLM-1. Permutation tests based on threshold-free-cluster-enhancement (TFCE) statistic.

| **Probability estimates $P_t$ (negative correlation)** | | | | |
|---|---|---|---|---|
| **Cluster** | **Hemisphere** | **Cluster size** | $p_{max}$ | **1-pmax(x,y,z)** |
| Occipital Pole | R | 62107 | 0 | (18,-98,22) |
| Frontal Pole | L | 676 | 0.036 | (-32,40,34) |
| **Belief revision $\Delta P_t$ (positive correlation)** | | | | |
| Frontal Orbital Cortex | L | 75881 | 0.002 | (-22,14,-16) |
| Frontal Pole | R | 424 | 0.045 | (34,36,32) |
| Supramarginal Gyrus, posterior division | L | 53 | 0.048 | (-68,-46,10) |

**Table S3.** Experiment 1: Probability estimates $(P_t)$ and belief revision $(\Delta P_t)$ contrasts based on GLM-1. Permutation tests based on cluster extent.

| **Probability estimates $P_t$ (negative correlation)** | | | | |
|---|---|---|---|---|
| **Cluster** | **Hemisphere** | **Cluster size** | $p_{max}$ | **1-$p_{max}$(x,y,z)** |
| Lateral Occipital Cortex, inferior division | R | 5326 | 0 | (56,-66,-12) |
| Temporal Fusiform Cortex, anterior division | L | 1989 | 0.001 | (-38,-8,-32) |
| Planum Polare | R | 926 | 0.003 | (52,6,-4) |
| Middle Temporal Gyrus, anterior division | L | 777 | 0.004 | (-52,-4,-18) |
| Postcentral Gyrus | R | 570 | 0.006 | (44,-18,36) |
| Right Cerebral White Matter | R | 394 | 0.009 | (36,0,-30) |
| Cingulate Gyrus, anterior division | R | 156 | 0.027 | (2,30,16) |
| Middle Temporal Gyrus, posterior division | L | 89 | 0.044 | (-62,-36,0) |
| Precuneus Cortex | - | 80 | 0.048 | (0,-56,56) |
| **Belief revision $\Delta P_t$ (positive correlation)** | | | | |
| Cingulate Gyrus, anterior division | L | 3082 | 0.002 | (-4,44,4) |
| Frontal Orbital Cortex | L | 2083 | 0.002 | (-24,14,-26) |
| Lingual Gyrus | R | 1274 | 0.004 | (2,-78,0) |
| Frontal Orbital Cortex | R | 708 | 0.008 | (14,12,-18) |
| Planum Polare | R | 582 | 0.01 | (44,-2,-20) |
| Brain-Stem | L | 566 | 0.01 | (-4,-36,-28) |
| Postcentral Gyrus | L | 390 | 0.014 | (-58,-20,24) |
| Supramarginal Gyrus, anterior division | R | 366 | 0.014 | (70,-20,24) |
| Lateral Occipital Cortex, superior division | L | 233 | 0.024 | (-32,-78,30) |

| | | | | |
|---|---|---|---|---|
| Precentral Gyrus | L | 207 | 0.028 | (-28,-8,54) |
| Precentral Gyrus | L | 188 | 0.03 | (-54,6,24) |
| Precentral Gyrus | R | 168 | 0.035 | (56,8,24) |

**Table S4.** Experiment 3: Instructed number $(IN_t)$ and difference in instructed number $(\Delta IN_t)$ contrasts based on GLM-1. For Experiment 3, $IN_t$ represents the two-digit number subjects were instructed to press at each period, and $\Delta IN_t$ represents the difference in number between successive periods. $IN_t$ is the control for $P_t$ in Experiment 1, and $\Delta IN_t$ is the control for $\Delta P_t$ in Experiment 1. Cluster-level inference using Gaussian random field theory (familywise error corrected at $p < .05$ with a cluster-forming threshold $z > 3.1$).

| **Instructed number $IN_t$ (negative correlation)** | | | | |
|---|---|---|---|---|
| **Cluster** | **Hemisphere** | **Cluster size** | **z-max** | **z-max(x,y,z)** |
| Precentral Gyrus | R | 1320 | 4.66 | (38,-22,62) |
| Postcentral Gyrus | L | 521 | 4.08 | (-40,-36,62) |
| Brain-Stem | L | 419 | 4.22 | (-4,-28,-10) |
| Cerebellar Left V | L | 401 | 4.8 | (-16,-50,-20) |
| Middle Frontal Gyrus | L | 316 | 4.22 | (-40,36,32) |
| Precentral Gyrus | R | 294 | 3.95 | (56,2,38) |
| Right Thalamus | R | 248 | 4.31 | (8,-18,0) |
| Superior Frontal Gyrus | L | 223 | 3.87 | (-20,-8,70) |
| **Difference in instructed number $\Delta IN_t$ (positive correlation)** | | | | |
| Precentral Gyrus | L | 1489 | 5.09 | (-40,-20,54) |

**Table S5.** Experiment 1 − Experiment 2 based on the probability estimates $(P_t)$ contrast in GLM-1. Cluster-level inference using Gaussian random field theory (familywise error corrected at $p < .05$ with a cluster-forming threshold $z > 3.1$).

| **Experiment 1 > Experiment 2 on negative probability estimates contrast** | | | | |
|---|---|---|---|---|
| **Cluster** | **Hemisphere** | **Cluster size** | **z-max** | **z-max(x,y,z)** |
| Paracingulate Gyrus | L | 7726 | 4.93 | (-8,38,-8) |
| Lateral Occipital Cortex, superior division | L | 3457 | 4.95 | (-16,-86,46) |
| Postcentral Gyrus | R | 328 | 3.82 | (48,-20,64) |

**Table S6.** Experiment 1 − Experiment 3: based on the probability estimates ($P_t$) contrast in GLM-1. Cluster-level inference using Gaussian random field theory (familywise error corrected at p<0.05 with a cluster-forming threshold z>3.1).

| **Experiment 1 > Experiment 2 on negative probability estimates contrast** | | | | |
|---|---|---|---|---|
| **Cluster** | **Hemisphere** | **Cluster size** | **z-max** | **z-max(x,y,z)** |
| Superior Temporal Gyrus, anterior division | L | 4099 | 5.04 | (-60,2,-2) |
| Occipital Pole | R | 2656 | 5.21 | (16,-98,28) |
| Central Opercular Cortex | R | 1575 | 5.41 | (62,-6,8) |
| **Experiment 1 > Experiment 2 on positive probability estimates contrast** | | | | |
| Supramarginal Gyrus, anterior division | L | 328 | 4 | (-52,-38,44) |

**Table S7.** Experiment 1: GLM-2. Cluster-level inference using Gaussian random field theory (familywise error corrected at $p < .05$ with a cluster-forming threshold $z > 3.1$).

| **Probability estimates $P_t$ (negative correlation)** | | | | |
|---|---|---|---|---|
| **Cluster** | **Hemisphere** | **Cluster size** | **z-max** | $z - \max(x, y, z)$ |
| Lingual Gyrus | L | 11504 | 6.2 | (-14,-52,-12) |
| (Local maxima) | | | | |
| Temporal fusiform cortex, posterior division | | | 5.12 | (−20, −46, −24) |
| Left accumbens | | | 5.11 | (−4, 8, −4) |
| Occipital pole | | | 5.11 | (16, −98, 28) |
| Superior temporal Gyrus | | | 5.10 | (−54, −4, −6) |
| Occipital pole | | | 5.06 | (12, −88, 40) |
| Central Opercular Cortex | R | 9872 | 6.19 | (60,-8,6) |
| (Local maxima) | | | | |
| Central Opercular Cortex | | | 6.13 | (56, −8, 10) |
| Central Opercular Cortex | | | 6.11 | (62, −4, 8) |
| Insular Cortex | | | 5.85 | (34, −14, 14) |
| Postcentral Gyrus | | | 5.85 | (44, −28, 64) |
| Precentral Gyrus | | | 5.54 | (40, −24, 58) |
| **Probability estimates (positive correlation)** | | | | |
| Postcentral Gyrus | L | 1680 | 5.45 | (-46,-26,56) |
| **Belief revision $\Delta P_t$ (positive correlation)** | | | | |
| Left Cerebral White Matter | L | 717 | 4.33 | (-20,18,-12) |
| Cingulate Gyrus, anterior division | - | 566 | 4.39 | (0,32,-8) |
| Frontal Orbital Cortex | R | 475 | 4.17 | (20,8,-16) |
| Postcentral Gyrus | L | 333 | 4.02 | (-30,-28,64) |
| **Intertemporal prior (negative correlation)** | | | | |
| Occipital Fusiform Gyrus | R | 1833 | 4.61 | (36,-74,-16) |
| Lateral Occipital Corte | R | 223 | 4.04 | (36,-66,-56) |
| Lateral Occipital Cortex | L | 204 | 4.25 | (-40,-82,-8) |

| $\ln(d)\times$ **signal (positive correlation)** | | | | |
|---|---|---|---|---|
| Middle Frontal Gyrus | R | 988 | 4.17 | (36,10,30) |
| Superior Frontal Gyrus | R | 821 | 4.56 | (6,34,48) |
| Superior Parietal Lobule | L | 620 | 4.26 | (-40,-52,56) |
| Supramarginal Gyrus | R | 604 | 4.29 | (46,-40,48) |
| Middle Frontal Gyrus | L | 277 | 4.21 | (-46,34,30) |

**Table S8** Experiment 1: GLM-2. Permutation tests based on the threshold-free-cluster-enhancement (TFCE) statistic.

| **Probability estimates $P_t$ (negative correlation)** | | | | |
|---|---|---|---|---|
| **Cluster** | **Hemisphere** | **Cluster size** | $p_{max}$ | $1 - p_{max}(x, y, z)$ |
| Postcentral Gyrus | R | 31490 | 0 | (38,-24,50) |
| Frontal Pole | R | 165 | 0.038 | (16,36,-20) |
| **Probability estimates $P_t$ (positive correlation)** | | | | |
| Postcentral Gyrus | L | 439 | 0.012 | (-42,-28,54) |
| **Belief revision $\Delta P_t$ (positive correlation)** | | | | |
| Right Caudate | R | 2857 | 0.027 | (12,14,-4) |
| Cingulate Gyrus, anterior division | R | 1873 | 0.027 | (4,26,24) |
| Postcentral Gyrus | L | 707 | 0.034 | (-42,-22,54) |
| Insular Cortex | L | 427 | 0.041 | (-42,6,-4) |
| Frontal Medial Cortex | L | 320 | 0.04 | (-4,40,-14) |
| Frontal Pole | L | 40 | 0.048 | (-2,60,28) |
| Insular Cortex | R | 20 | 0.048 | (40,6,-14) |
| **Intertemporal prior (negative correlation)** | | | | |
| Lateral Occipital Cortex, inferior division | R | 2117 | 0.006 | (38,-74,14) |
| Frontal Pole | R | 185 | 0.038 | (36,42,36) |
| Frontal Pole | R | 114 | 0.043 | (40,44,-6) |
| Middle Frontal Gyrus | R | 14 | 0.048 | (34,28,50) |
| Right Thalamus | R | 13 | 0.046 | (22,-28,-2) |
| Frontal Pole | R | 1 | 0.05 | (32,48,-8) |
| Frontal Pole | R | 1 | 0.05 | (30,52,-6) |

**Table S9.** Experiment 1: GLM-2. Permutation tests based on cluster extent.

| **Probability estimates $P_t$ (negative correlation)** | | | | |
|---|---|---|---|---|
| **Cluster** | **Hemisphere** | **Cluster size** | $p_{max}$ | $1 - p_{max}(x, y, z)$ |
| Postcentral Gyrus | R | 1770 | 0.001 | (42,-20,38) |
| Lateral Occipital Cortex, inferior division | R | 1085 | 0.003 | (60,-68,-6) |
| Lateral Occipital Cortex, inferior division | L | 674 | 0.006 | (-42,-84,10) |
| Temporal Fusiform Cortex, posterior division | L | 597 | 0.007 | (-36,-18,-32) |
| Heschl's Gyrus | R | 527 | 0.008 | (54,-12,2) |
| Temporal Fusiform Cortex, anterior division | R | 441 | 0.01 | (34,-2,-30) |
| Cerebellar Left V | L | 235 | 0.022 | (-16,-50,-24) |
| Middle Temporal Gyrus, anterior division | L | 193 | 0.027 | (-52,-4-18) |
| Frontal Medial Cortex | L | 134 | 0.039 | (-10,34,-18) |
| Lingual Gyrus | L | 134 | 0.039 | (-6,-48,-2) |
| **Probability estimates $P_t$ (positive correlation)** | | | | |
| Supramarginal Gyrus, anterior division | L | 1012 | 0.002 | (-44,-30,40) |
| **Belief revision $\Delta P_t$ (positive correlation)** | | | | |
| Cingulate Gyrus, anterior division | - | 450 | 0.016 | (0,34,14) |
| Frontal Orbital Cortex | R | 351 | 0.021 | (16,12,-16) |
| Frontal Orbital Cortex | L | 300 | 0.024 | (-22,10,-18) |
| Postcentral Gyrus | L | 215 | 0.035 | (-38,-24,46) |
| **Intertemporal prior (negative correlation)** | | | | |
| Temporal Occipital Fusiform Cortex | R | 1687 | 0.002 | (34,-60,24) |

| | | | | |
|---|---|---|---|---|
| Lateral Occipital Cortex, superior division | R | 220 | 0.042 | (38,-70,32) |
| **$\ln(d)\times$ signal (positive correlation)** | | | | |
| Superior Parietal Lobule | L | 384 | 0.022 | (-38,-52,40) |

**Table S10.** Model-fitting summary.

| **Model** | **Likelihood** | **Number of Parameters** | **AIC** |
|---|---|---|---|
| SN-original | $356.27 \pm 44.7$ | 6 | $-700.54 \pm 89.49$ |
| SN-SigDep-$\beta$ | $383.08 \pm 44.69$ | 9 | $-748.15 \pm 89.37$ |
| SN-SigDep-$\alpha$ | $398.61 \pm 45.78$ | 9 | $-779.21 \pm 91.57$ |
| SN-SigDep-$\alpha\beta$ | $410.03 \pm 46.37$ | 12 | $-796.07 \pm 92.74$ |

**Table S11.** Model comparison using paired t-test with Bonferroni correction.

| Model | $t$ test ($df = 29$) | $p$-value | Bonferroni-corrected $p$-value |
|---|---|---|---|
| SN-SigDep-$\beta$ $-$ SN-original | 5.15 | 0.0000 | 0.0001 |
| SN-SigDep-$\alpha$ $-$ SN-original | 6.93 | 0.0000 | 0.0000 |
| SN-SigDep-$\alpha\beta$ $-$ SN-original | 8.56 | 0.0000 | 0.0000 |
| SN-SigDep-$\alpha$ $-$ SN-SigDep-$\beta$ | 4.01 | 0.0004 | 0.0023 |
| SN-SigDep-$\alpha\beta$ $-$ SN-SigDep-$\beta$ | 6.55 | 0.0000 | 0.0000 |
| SN-SigDep-$\alpha\beta$ $-$ SN-SigDep-$\alpha$ | 5.03 | 0.0000 | 0.0001 |

**Table S12.** Experiment 1: GLM-1 without the action-handedness regressor. Cluster-level inference using Gaussian random field theory (familywise error corrected at $p < .05$ with a cluster-forming threshold $z > 3.1$).

| **Probability estimates $P_t$ (negative correlation)** | | | | |
|---|---|---|---|---|
| **Cluster** | **Hemisphere** | **Cluster size** | **z-max** | **z-max(x,y,z)** |
| Temporal Occipital Fusiform Cortex | L | 27876 | 6.63 | (-24,-54,-22) |
| Supplementary Motor Cortex | R | 1096 | 4.72 | (12,0,46) |
| **Probability estimates $P_t$ (positive correlation)** | | | | |
| Postcentral Gyrus | L | 705 | 4.73 | (-46,-28,54) |
| **Belief revision $\Delta P_t$ (positive correlation)** | | | | |
| Frontal Orbital Cortex | L | 8939 | 5.02 | (-24,20,-10) |
| Frontal Orbital Cortex | R | 2244 | 4.7 | (20,6,-14) |
| Lateral Occipital Cortex, superior division | L | 1786 | 4.45 | (-28,-82,34) |
| Postcentral Gyrus | L | 1752 | 4.61 | (-58,-24,38) |
| Postcentral Gyrus | R | 1053 | 4.61 | (60,-18,36) |
| Precentral Gyrus | L | 361 | 3.92 | (-52,6,32) |

**Table S13.** Experiment 1: GLM-2 without $P_t$ and $\Delta P_t$ regressors. Cluster-level inference using Gaussian random field theory (familywise error corrected at $p < .05$ with a cluster-forming threshold $z > 3.1$).

| **Intertemporal prior (negative correlation)** | | | | |
|---|---|---|---|---|
| **Cluster** | **Hemisphere** | **Cluster size** | **z-max** | $\boldsymbol{z-\max(x,y,z)}$ |
| Temporal Occipital Fusiform Cortex | R | 2101 | 5.3 | (44,-54,-20) |
| Lateral Occipital Cortex, inferior division | L | 976 | 4.49 | (-40,-82,-8) |
| Paracingulate Gyrus | R | 316 | 4.12 | (14,34,30) |
| Right Thalamus | R | 240 | 4.32 | (22,-28,-4) |
| Precuneus Cortex | R | 230 | 4.13 | (6,-82,48) |
| Frontal Pole | R | 186 | 3.73 | (42,44,34) |
| $\mathbf{ln}(\boldsymbol{d})\times$ **signal (positive correlation)** | | | | |
| Supramarginal Gyrus, posterior division | L | 945 | 4.58 | (-50,-42,56) |
| Supramarginal Gyrus, posterior division | R | 923 | 4.44 | (46,-40,48) |
| Middle Frontal Gyrus | R | 785 | 4.19 | (46,12,50) |
| Superior Frontal Gyrus | R | 521 | 4.49 | (6,32,50) |
| Middle Frontal Gyrus | L | 226 | 4.1 | (-46,34,30) |

### Task Instruction (in Chinese) for Experiment 1

All subjects were instructed in Chinese with this document. We also included the English translation version after the Chinese version.

#### 實驗介紹

**簡介：**

想像我們手邊有兩個箱子：紅箱子與藍箱子。紅箱子中有比較多的紅球；藍箱子中有比較多的藍球。我們會從箱子中抽球給您看，您會知道球的顏色，但不知道球是從紅箱子或藍箱子抽出的。您要做的作業為猜測該球是從藍箱子抽出的機率為何。

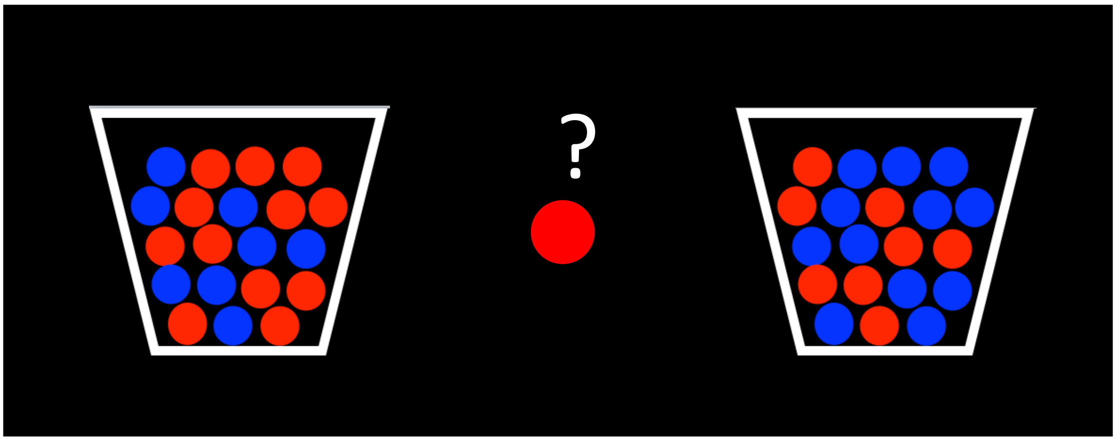


Color ratio= [60 40]　　Color ratio= [40 60]

抽球規則：

1. 一題會抽出10顆球。一次抽出一顆球。
2. 抽球並不會都從同一個箱子。一開始設定的箱子為紅箱子，但從抽第一顆球開始，電腦便有一定的機率轉換成由藍箱子抽球。

3. 箱子最多只會轉換一次。一旦轉換到藍箱子， 就不會再轉換回紅箱子。
4. 球取後會被放回原箱子。故在同一題中兩箱子個紅藍球比例會是固定的。且兩箱子的顏色比例是相反的。

**實驗流程**

首先，您會看到該題「箱子間轉換的機率」與「箱子中紅藍球的比例」

您會看到電腦從箱子中抽出一顆球，看到球後即可估計球是從藍箱子抽出的機率。每題共有10次觀察。

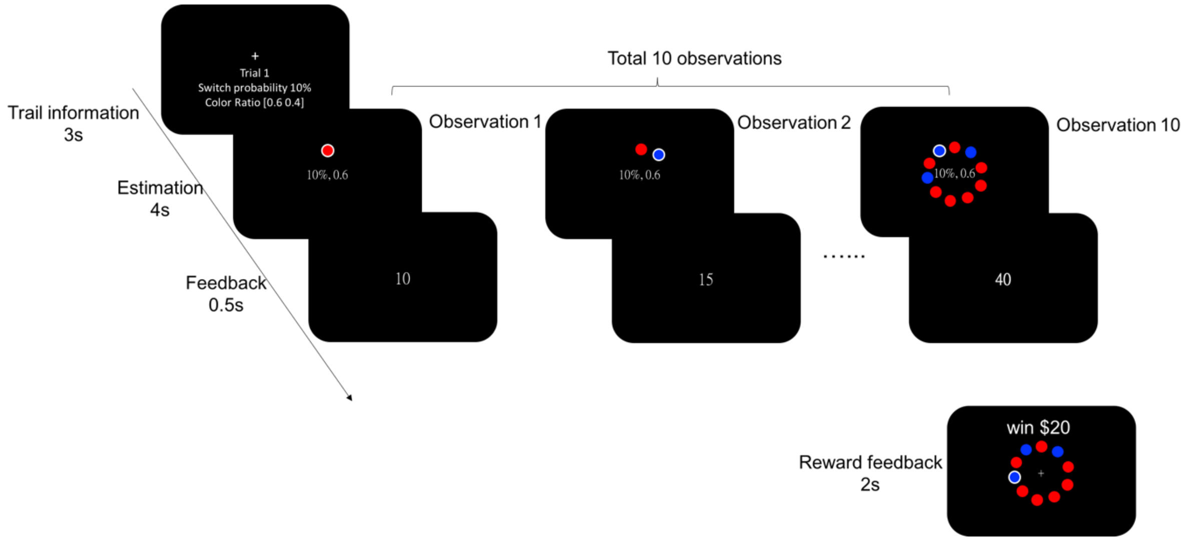


在實驗中，每題都會提供給您以下的資訊以幫助您做機率的估計，這些資訊在您做估計時都會提供給您：

1. 紅箱子中紅藍球的比例。(您可以依據這個比例知道藍箱子的比例)

2. 轉換機率：在每次觀察中，由紅箱子抽球轉換到由藍箱子抽球的機率。

**什麼是轉換機率？**

假設轉換機率為25%，那表示每次觀察電腦都有25%的機率由紅箱子轉換到由藍箱子抽球。這表示第一次觀察您所看到的球有25%的機率來自藍箱子，75%的機率來自紅箱子。一旦轉換成由藍箱子抽球，那麼直到這一題十次觀察結束，都會由藍箱子抽球。若第一次觀察到的球是由紅箱抽出，那麼下一次觀察的球是由藍箱子抽出的機率也會是25%。每次觀察的轉換機率為獨立。因此，轉換機率其實就是在每次觀察中，由紅箱子抽球轉換到由藍箱子抽球的機率。當然，電腦也有可能10次都由紅箱抽球，或10次都由藍箱抽球。

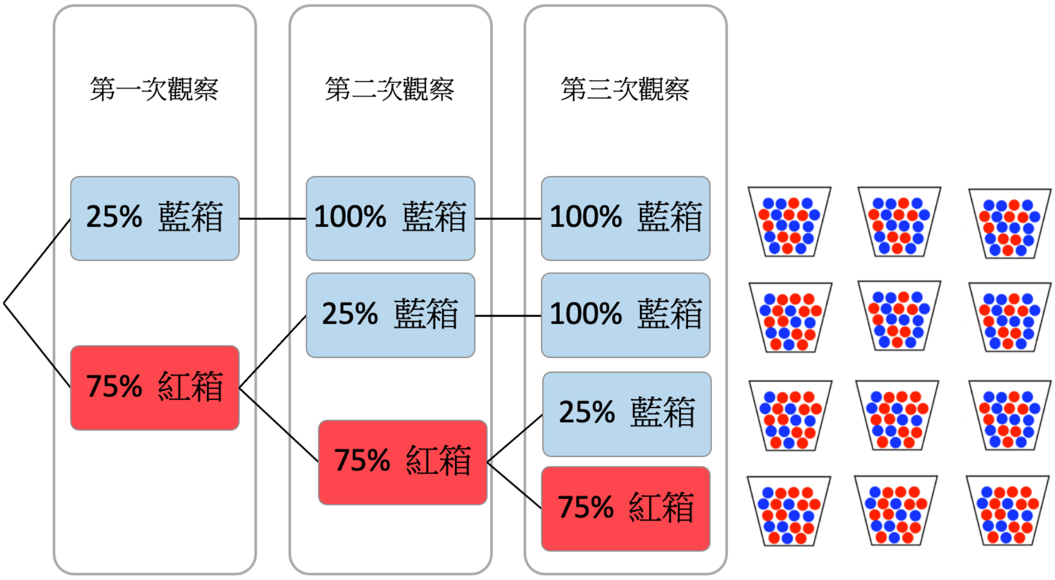

## 範例

酬賞: 一題（十次嘗試）最多可獲得30元，實驗結束時會抽十題實現。

計算方式$30\times (\$0.10 - (\$0.20 \times Error^2))$ 猜測越近可以得到越多錢

Period 1:

猜測球是從藍箱子抽出的機率為10%，實際上是由紅箱子抽出。

$30*(\$0.10 - (\$0.20 \times 0.1^2) = 2.94$

Period 7:

猜測球是從藍箱子抽出的機率為10%，實際上是由藍箱子抽出。

$30*(\$0.10 - (\$0.20 \times 0.9^2) = -1.86$

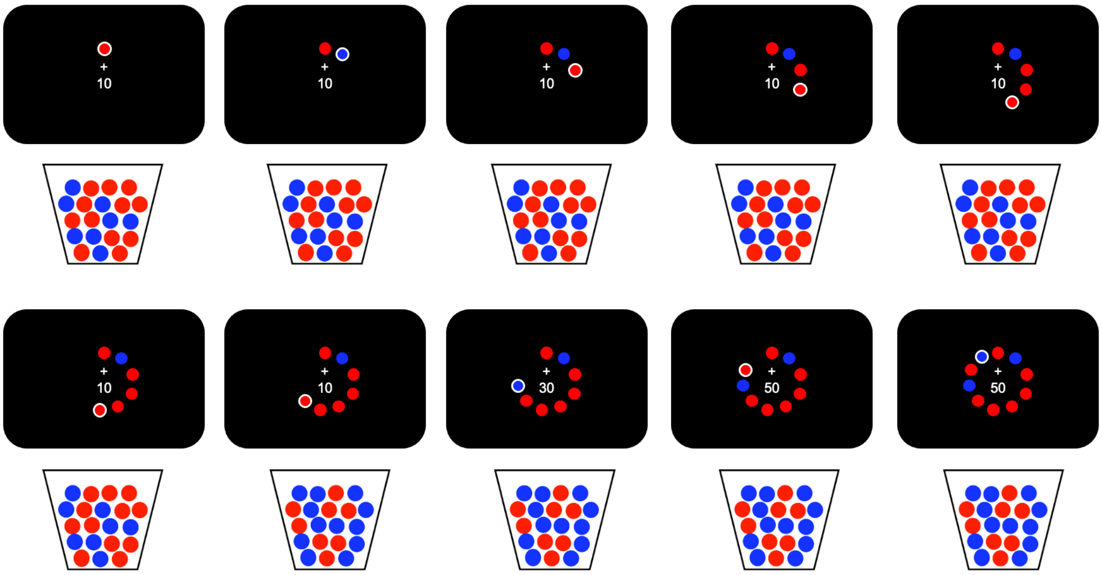

### Task Instruction for Experiment 1

**Brief Introduction:**

Imagine two boxes, red and blue, placed in front of you. In the red box, there are more red balls. In the blue box, there are more blue balls, as shown below. We will draw balls from one of the boxes and show them to you. You will see the color of the drawn balls, but you will not know whether they come from the red or blue box. Your task is to estimate the probability that the drawn ball comes from the blue box.

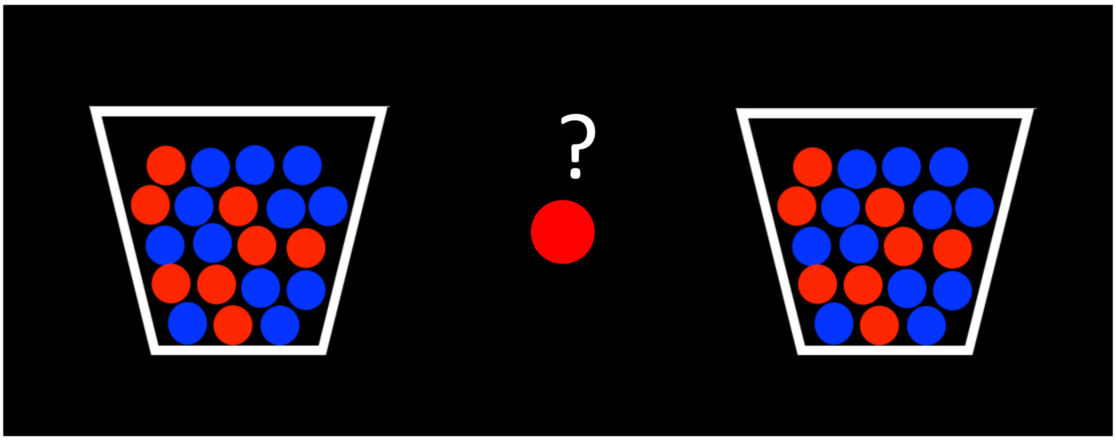


Color ratio= [60 40] Color ratio= [40 60]

Ball drawing rules：

1. On each trial, we will draw a total of 10 balls. We will draw one ball at a time.
2. We will not always draw from the same box. In the beginning, we will always start from the red box. But before we draw the first ball, there is a certain probability that the box we draw from would switch to the blue box.
3. A shift would happen at most one time during a trial. Once a shift to the blue box happens, it is no longer possible to shift again from the blue to the red box.
4. Once a ball is drawn and shown to you, it would be returned back to the box. Therefore, during a trial the ratio of blue and red balls in both boxes would be fixed. And the color ratio is opposite between the two boxes.

**Experiment Procedure**

At the beginning of a trial, you will first see information about the transition probability and the color ratio of this trial. Then, you will see a ball drawn from a box (an observation). When you see the ball, estimate the probability that it comes from the blue box. There are a total of 10 observations in a trial.

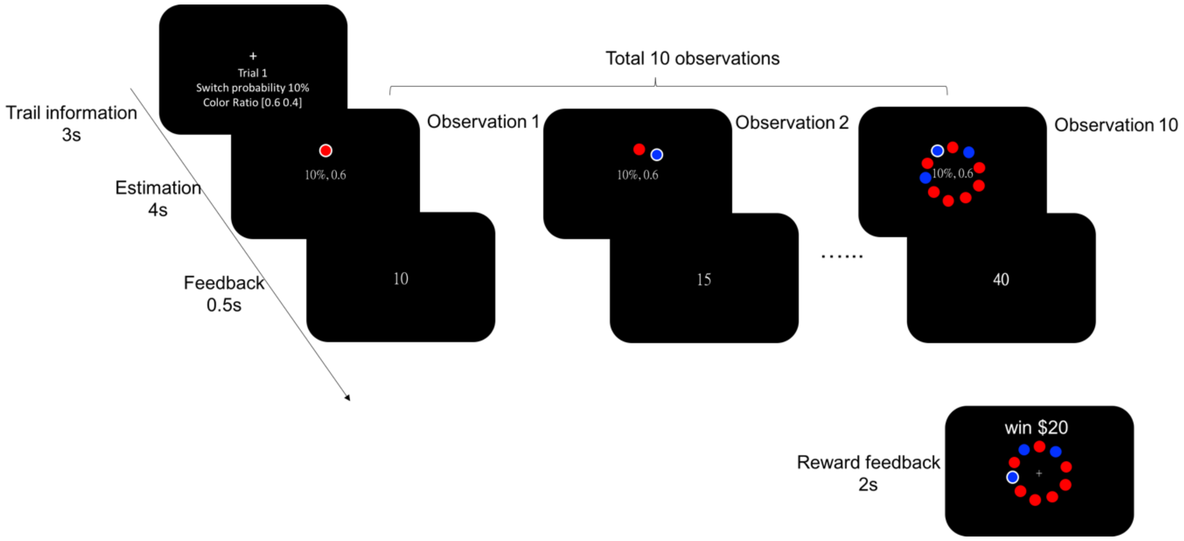


During the experiment, in each trial we will give you the following information to help you estimate probability:

1. The ratio of the red and blue balls in the red box (you can use this ratio to know the proportion of red and blue balls in the blue box).

2. Transition probability: this is the probability that, at each observation, we shift from drawing the ball from the red box to the blue box.

**What is transition probability?**

Suppose that transition probability is 25%. This indicates that at each observation there is a 25% chance that the box switches from the red box to the blue box. It means that at the first observation, there is a 25% chance that the drawn ball comes from the blue box, and a 75% chance that the ball comes from the red box. During a trial, once the box transitions from the red to the blue box, all the remaining balls will be drawn from the blue box. If the drawn ball at the first observation comes from the red box, then at the second observation the probability that the ball comes from the blue box would also be 25%. At each observation the transition probability is independent. In other words, transition probability is, at each observation, the probability that the box switches from the red to the blue box. Of course, it is possible that all the 10 observations come from the red box, or all 10 observations comes from the blue box.

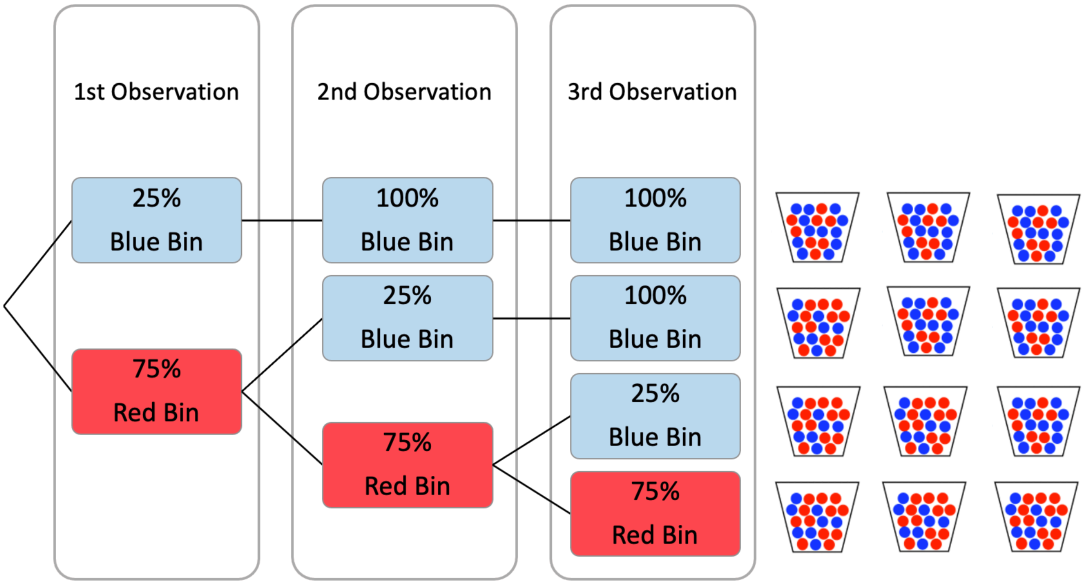

**Example**

Reward: you can earn at most 30 dollars in a trial (10 observations). At the end of the experiment, we will randomly select 10 trials to realize your payoffs.

Reward calculation formula: $30\times$ ($0.10 - ($0.20 $\times$ $Error^2$)). This means that when your estimate is closer to the true box where the ball is drawn you would receive more money.

Observation 1:

You guess that the probability that the ball comes from the blue box is 10%. However, it was in fact coming from the red box. As a result, you win 30*($0.10 - ($0.20 $\times$ $0.1^2$) = 2.94 at this observation.

Observation 7:

You guess that the probability that the ball comes from the blue box is 10%. However, it was in fact coming from the red box. As a result, you lose 30*($0.10 - ($0.20 $\times$ $0.9^2$) = -1.86 at this observation.

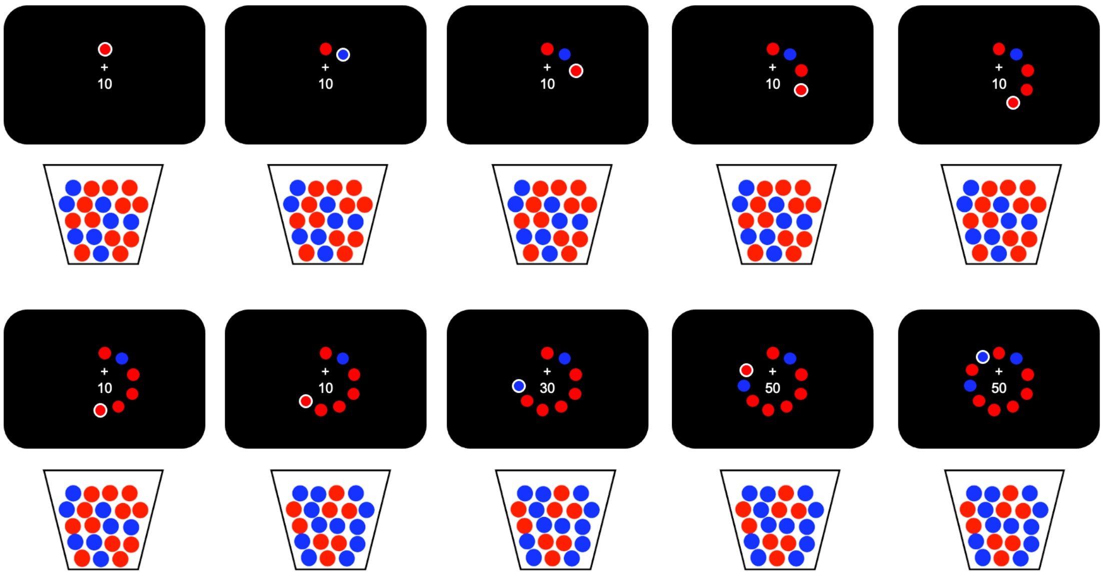

#### Demo of the trial sequence

Here we show the sequence of two example trials.

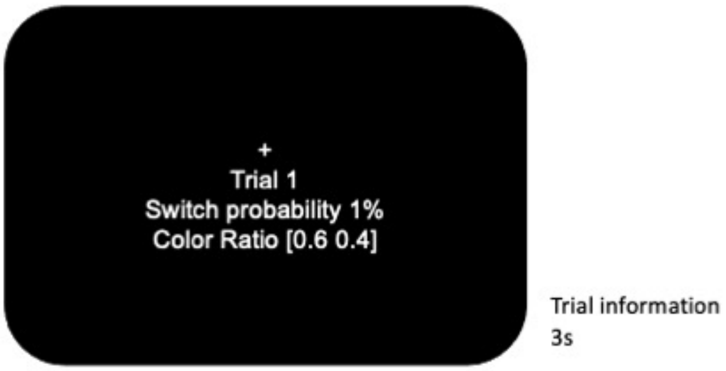


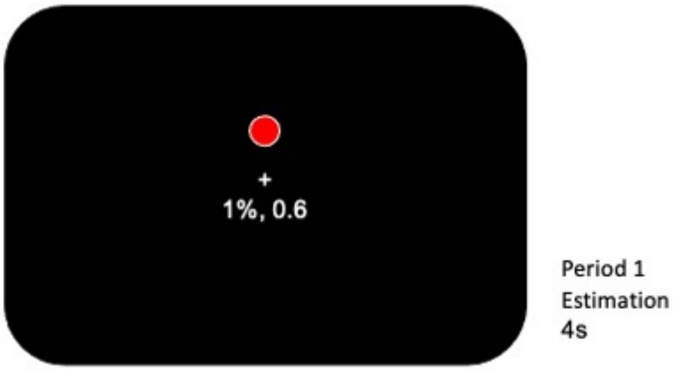

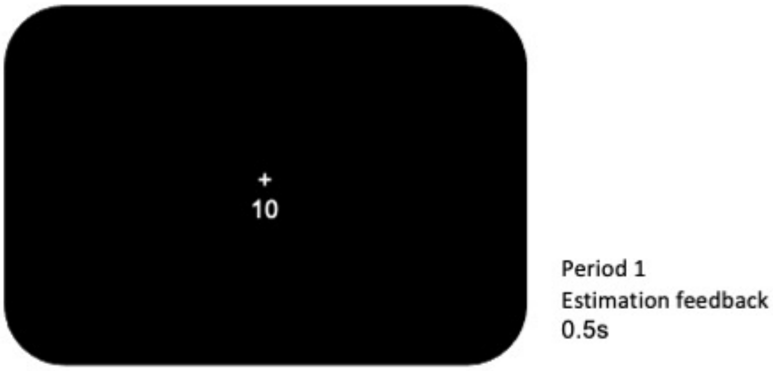
+
10
Period 1
Estimation feedback
0.5s

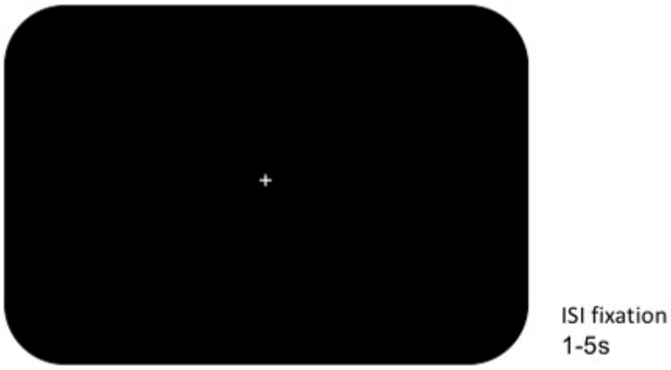
+
ISI fixation
1-5s

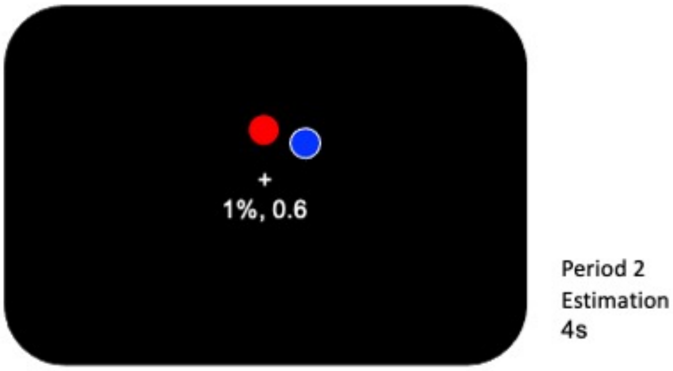
+
1%, 0.6
Period 2
Estimation
4s

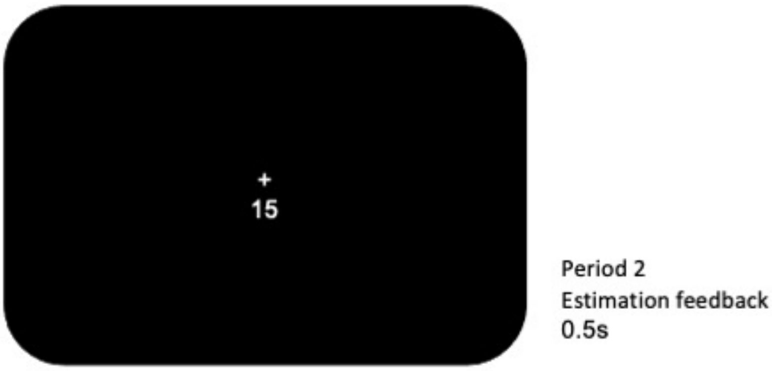
15
Period 2
Estimation feedback
0.5s

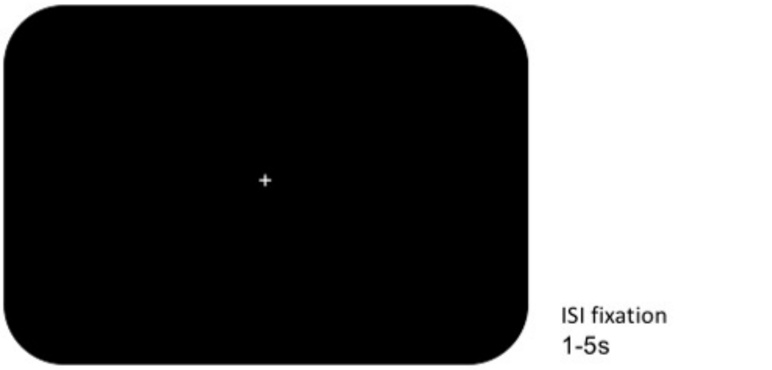
ISI fixation
1-5s

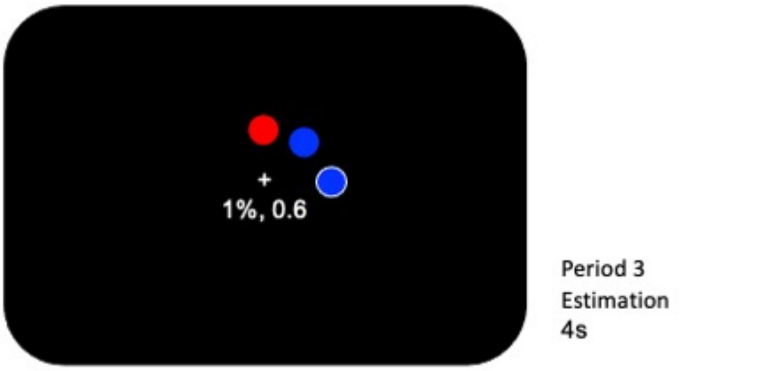
1%, 0.6
Period 3
Estimation
4s

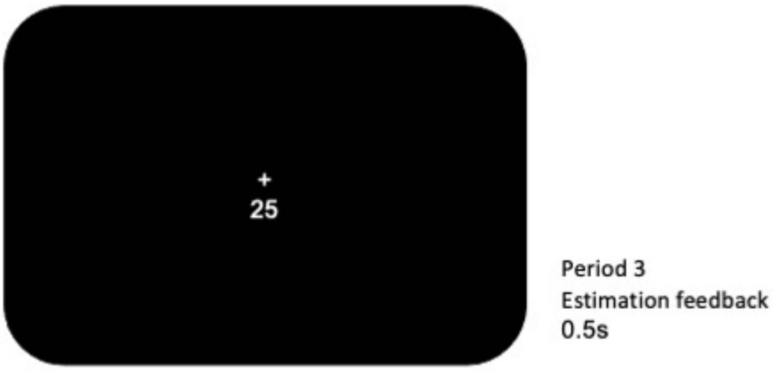
+
25
Period 3
Estimation feedback
0.5s

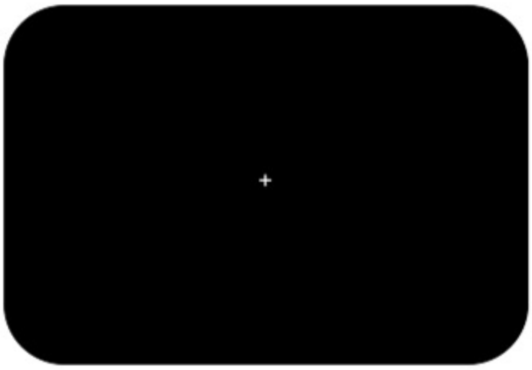

ISI fixation
1-5s

1

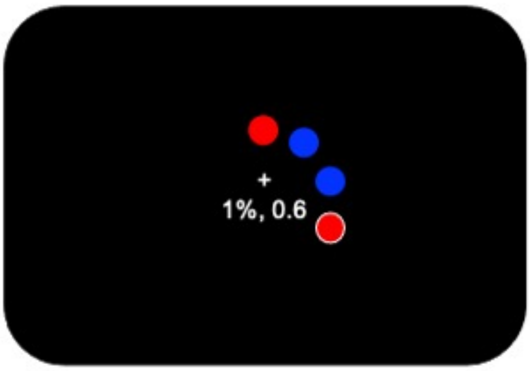


Period 4
Estimation
4s

2

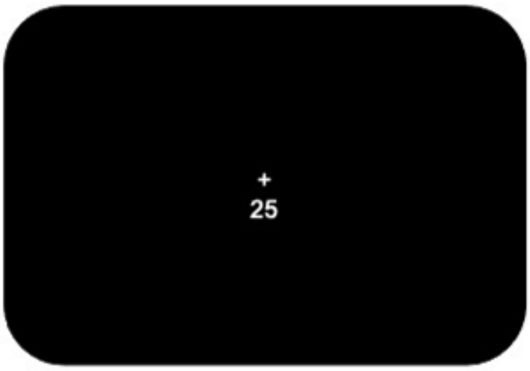


Period 4
Estimation feedback
0.5s

3

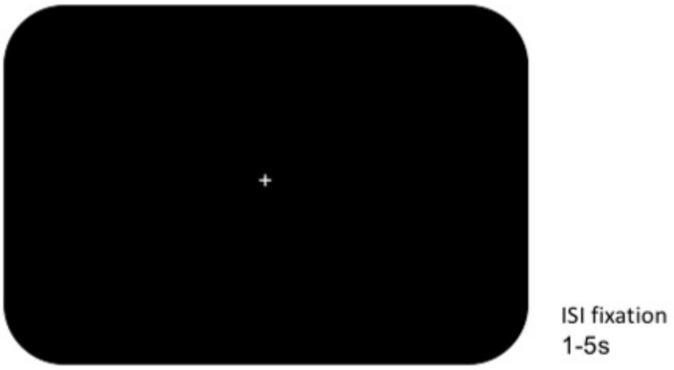
+
ISI fixation
1-5s

1

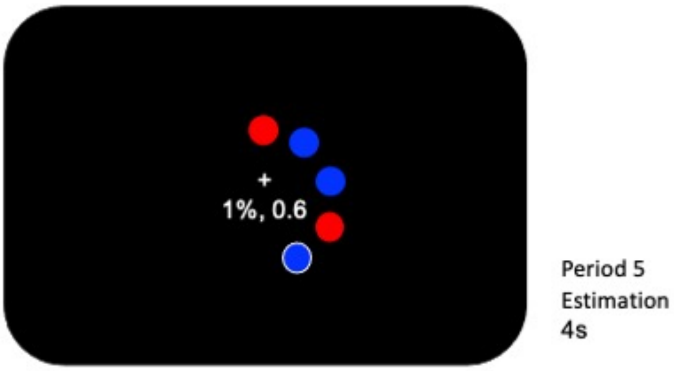
+
1%, 0.6
Period 5
Estimation
4s

2

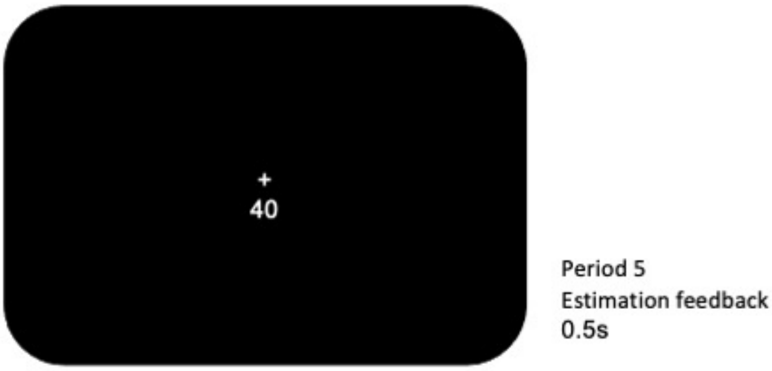
+
40
Period 5
Estimation feedback
0.5s

3

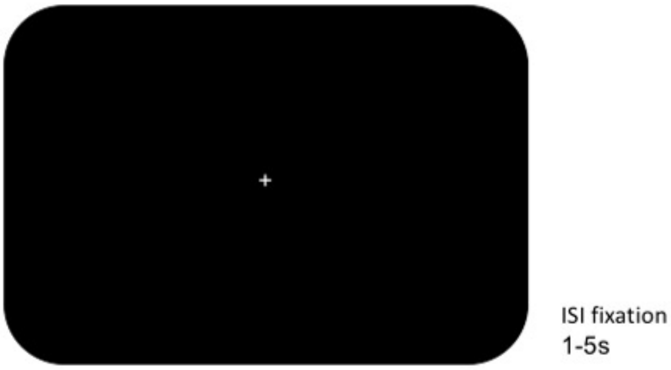
+
ISI fixation
1-5s

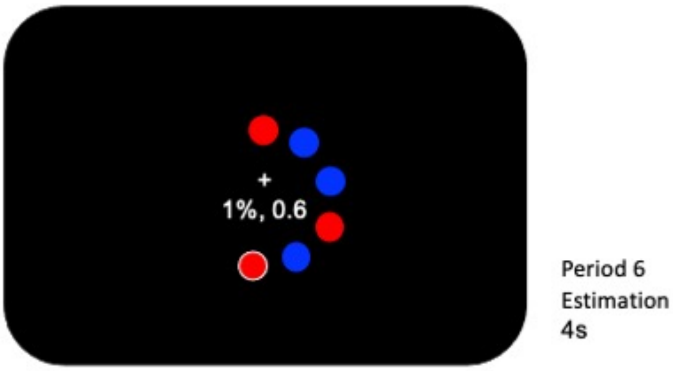
+
1%, 0.6
Period 6
Estimation
4s

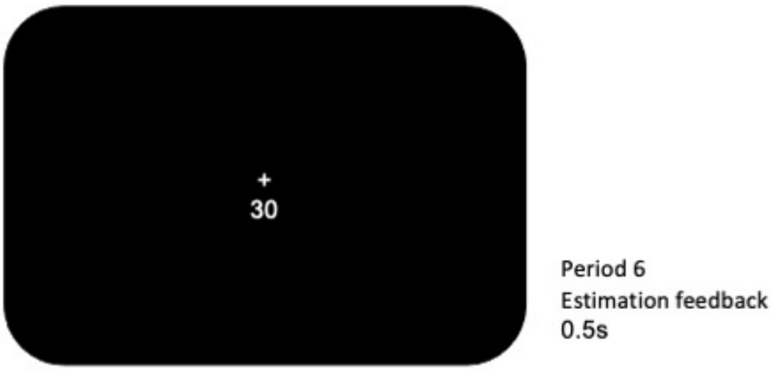
+
30
Period 6
Estimation feedback
0.5s

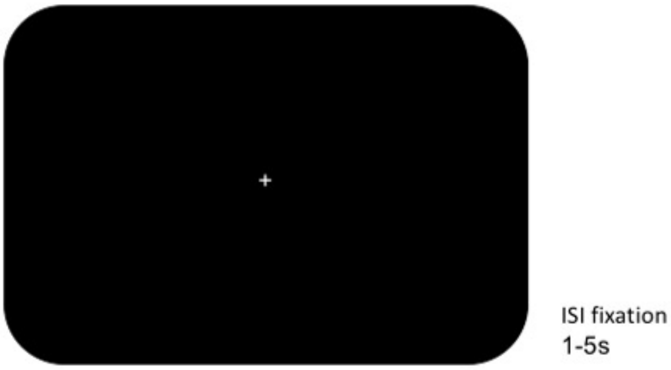
+
ISI fixation
1-5s

1

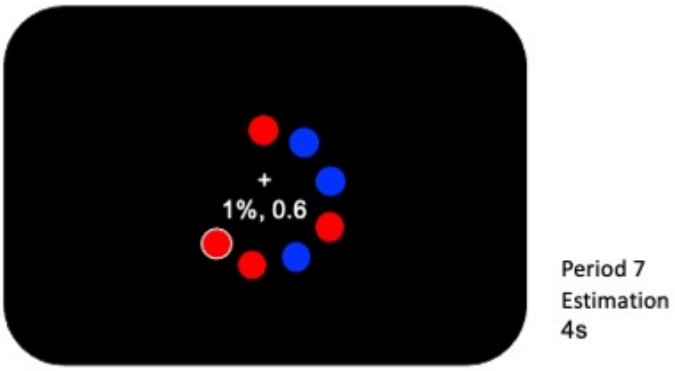
+
1%, 0.6
Period 7
Estimation
4s

2

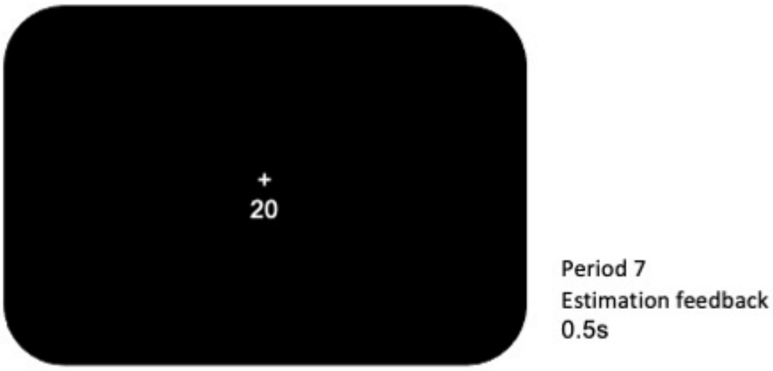
+
20
Period 7
Estimation feedback
0.5s

3

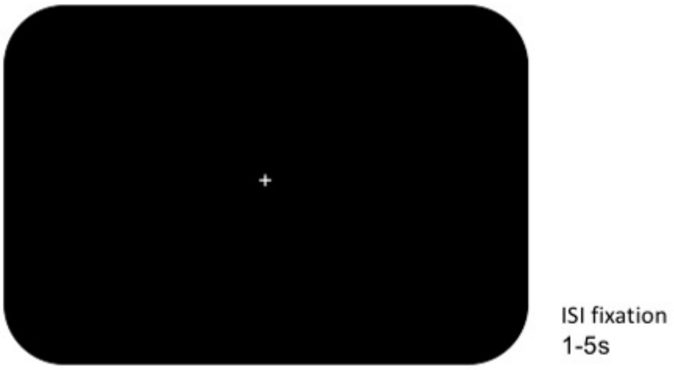
+
ISI fixation
1-5s

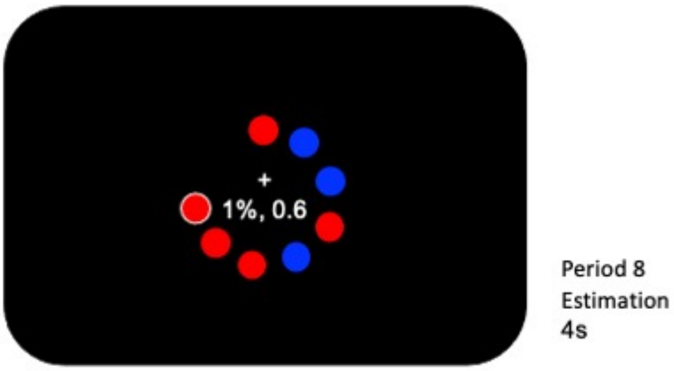
+
1%, 0.6
Period 8
Estimation
4s

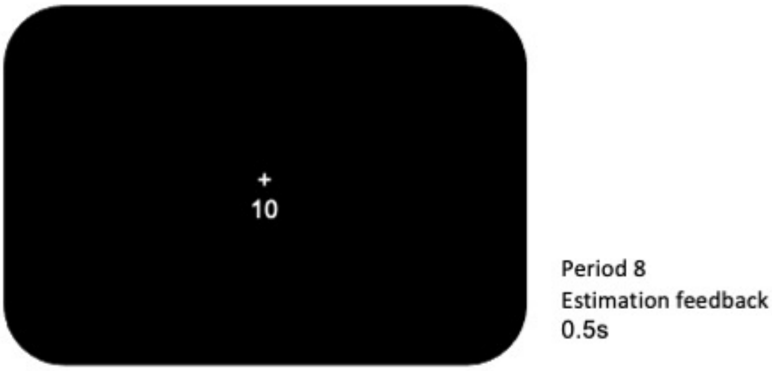
+
10
Period 8
Estimation feedback
0.5s

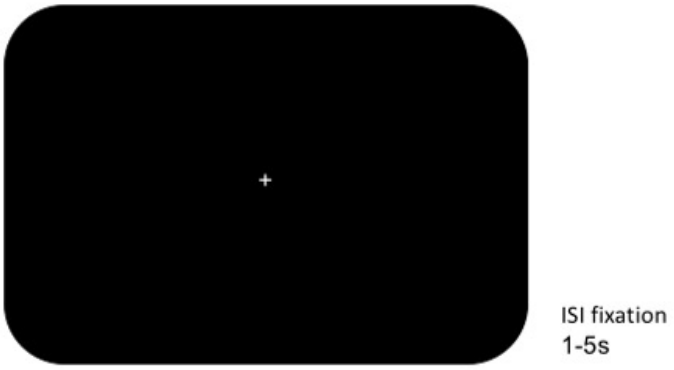
+
ISI fixation
1-5s

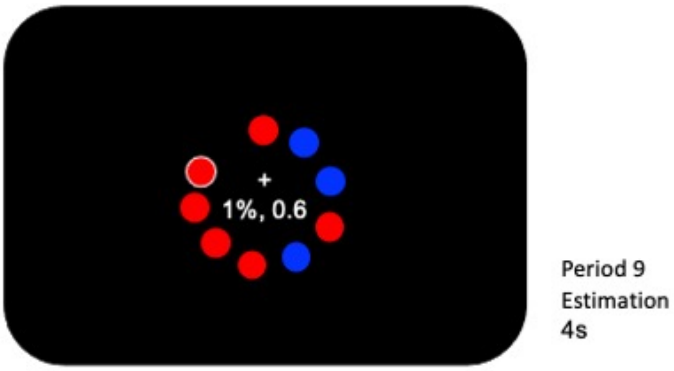
+
1%, 0.6
Period 9
Estimation
4s

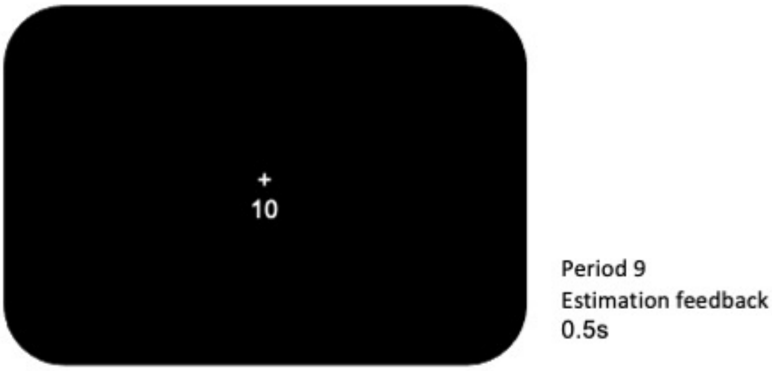
+
10
Period 9
Estimation feedback
0.5s

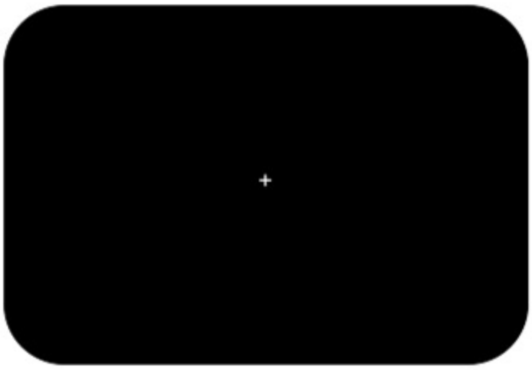


ISI fixation
1-5s

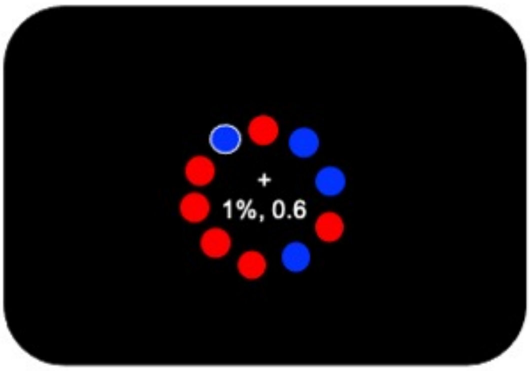


Period 10
Estimation
4s

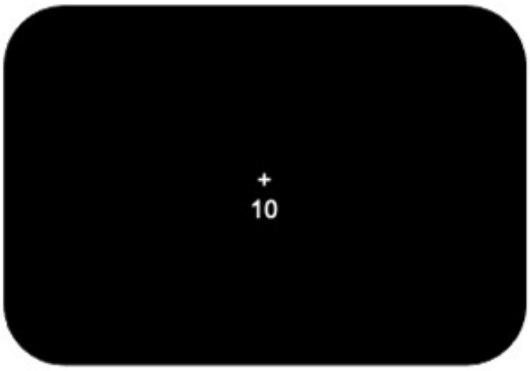


Period 10
Estimation feedback
0.5s

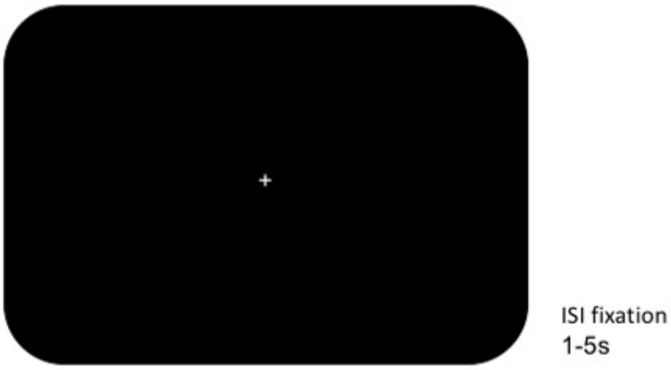


1

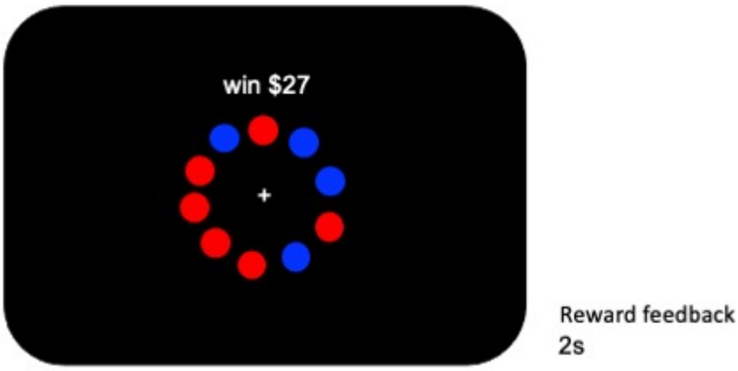


2

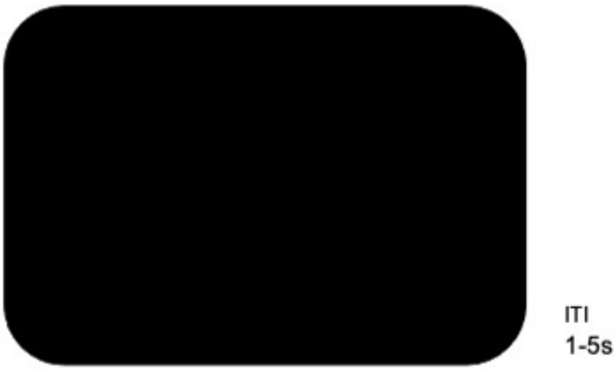


3

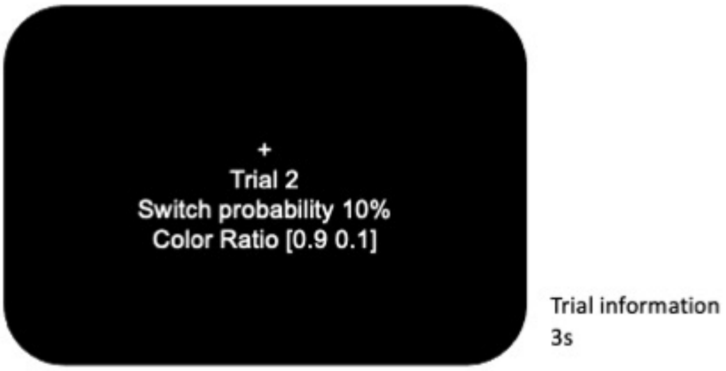
+
Trial 2
Switch probability 10%
Color Ratio [0.9 0.1]
Trial information
3s

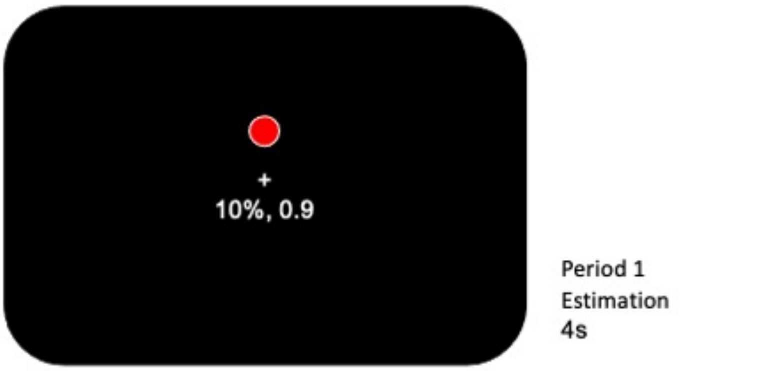
+
10%, 0.9
Period 1
Estimation
4s

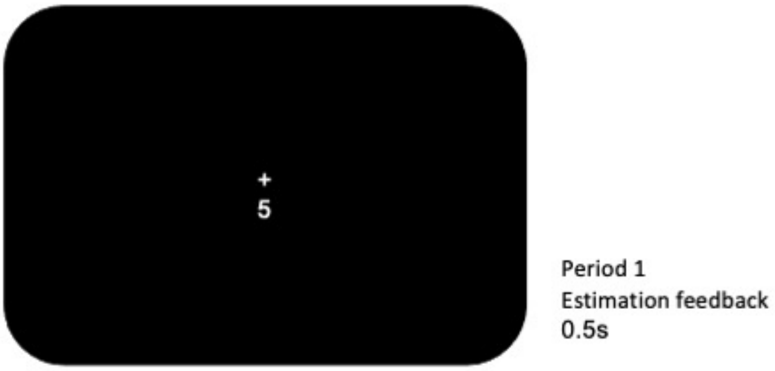
+
5
Period 1
Estimation feedback
0.5s

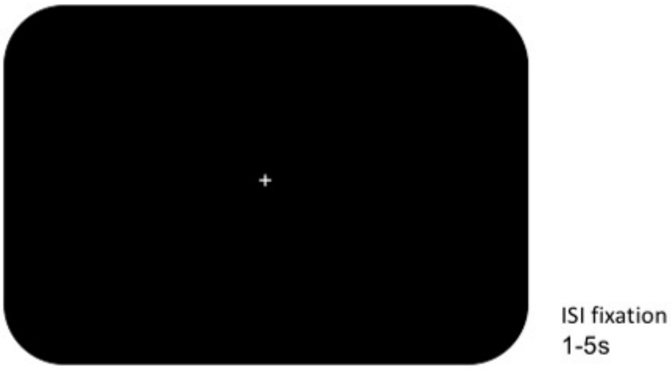
+
ISI fixation
1-5s

1

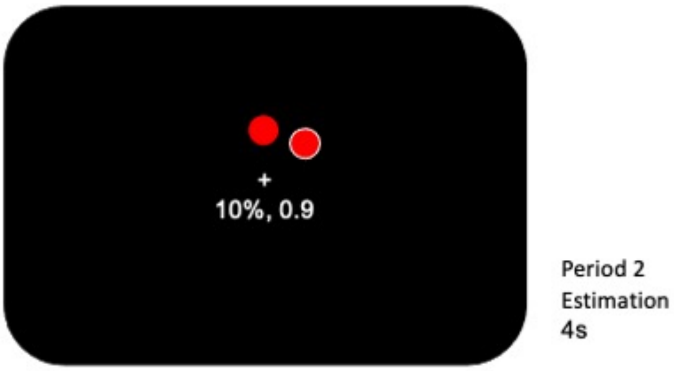
+
10%, 0.9
Period 2
Estimation
4s

2

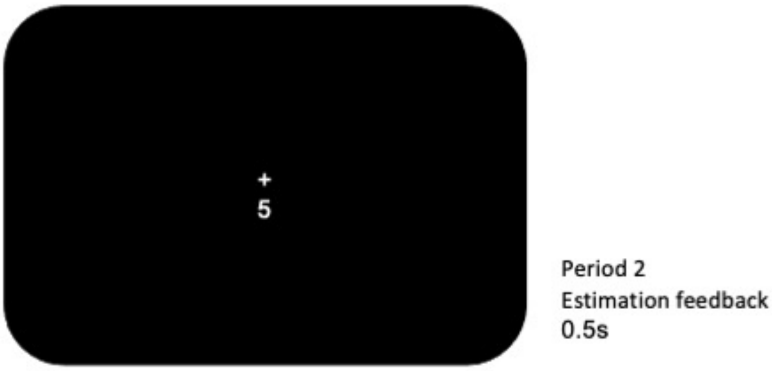
+
5
Period 2
Estimation feedback
0.5s

3

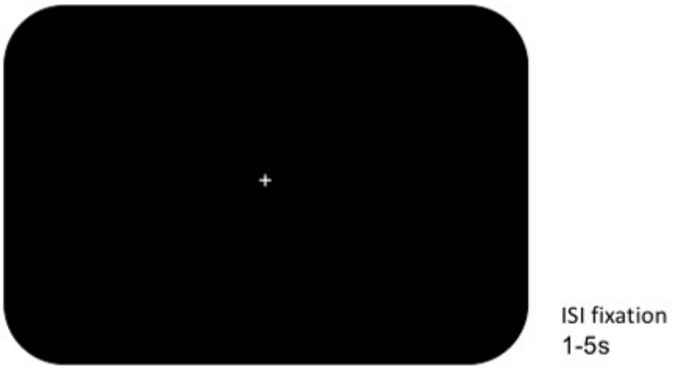
+
ISI fixation
1-5s

1

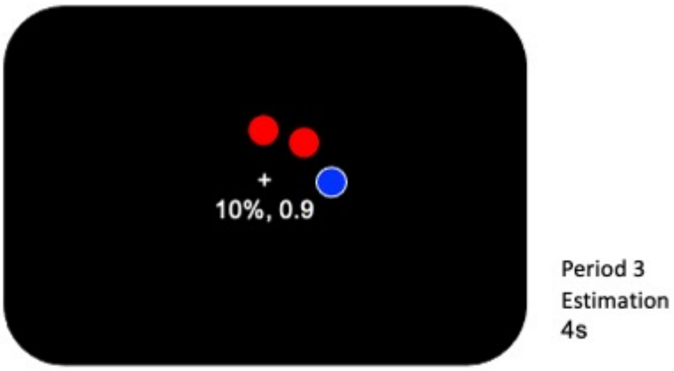
+
10%, 0.9
Period 3
Estimation
4s

2

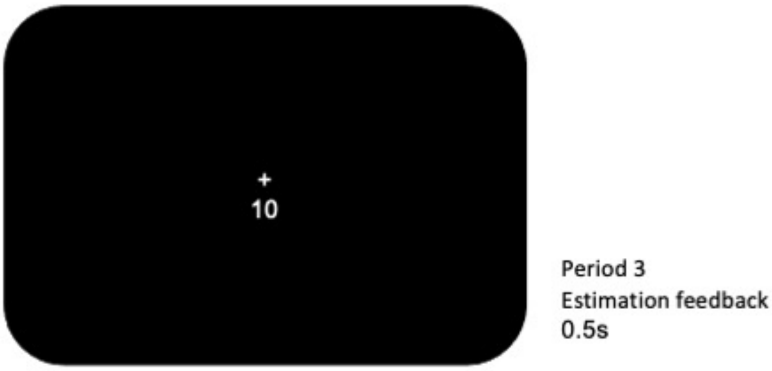
+
10
Period 3
Estimation feedback
0.5s

3

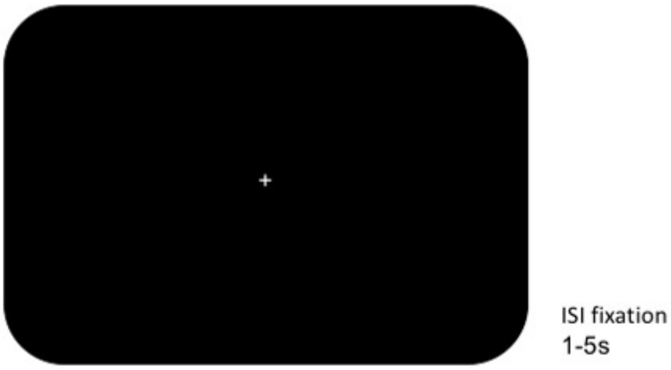


1

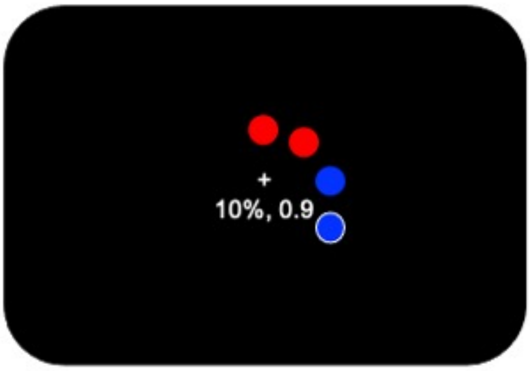


Period 4
Estimation
4s

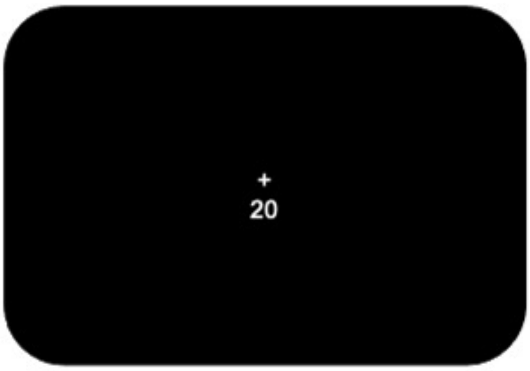


Period 4
Estimation feedback
0.5s

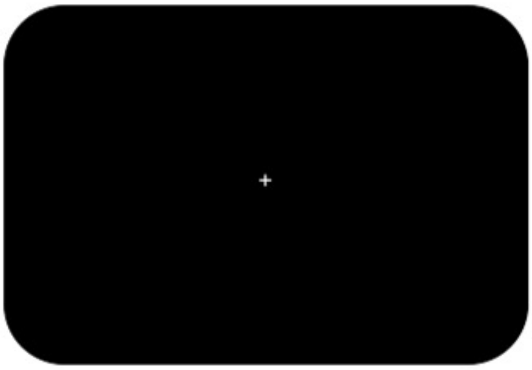

ISI fixation
1-5s

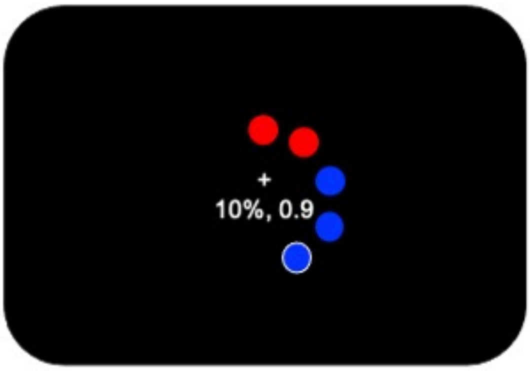


Period 5
Estimation
4s

1

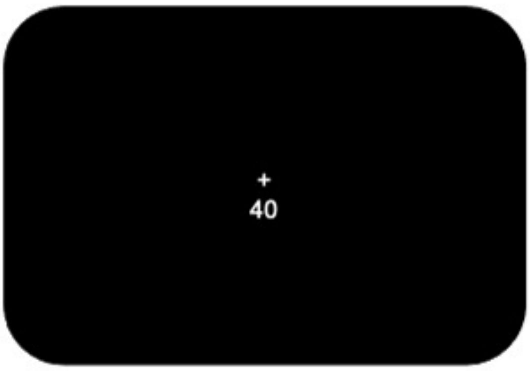


Period 5
Estimation feedback
0.5s

2

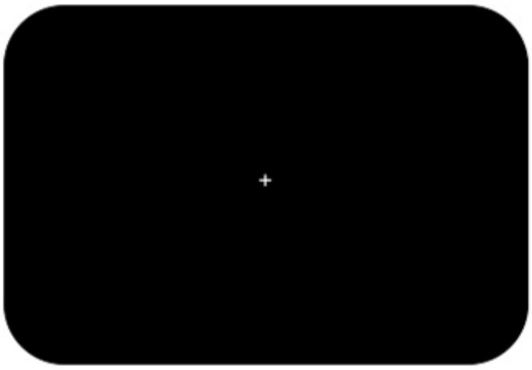

ISI fixation
1-5s

3

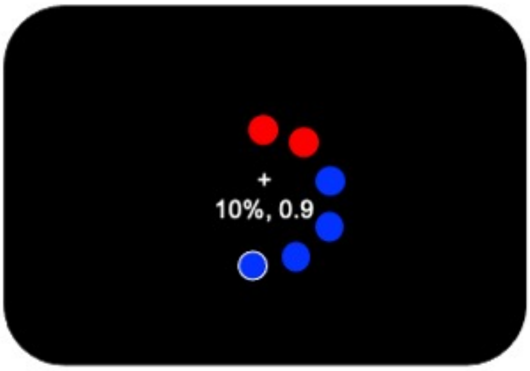


Period 6
Estimation
4s

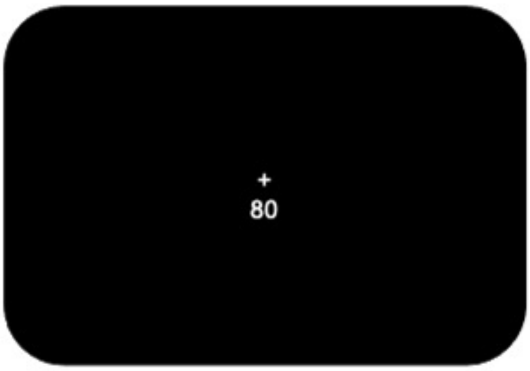


Period 6
Estimation feedback
0.5s

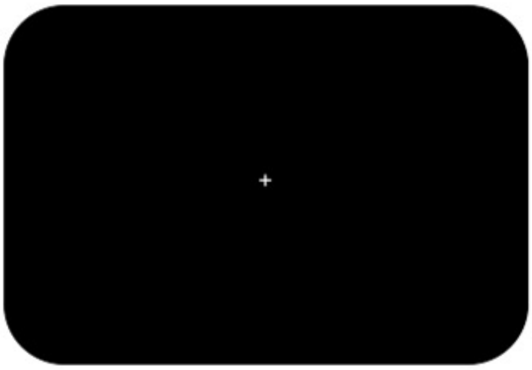


ISI fixation
1-5s

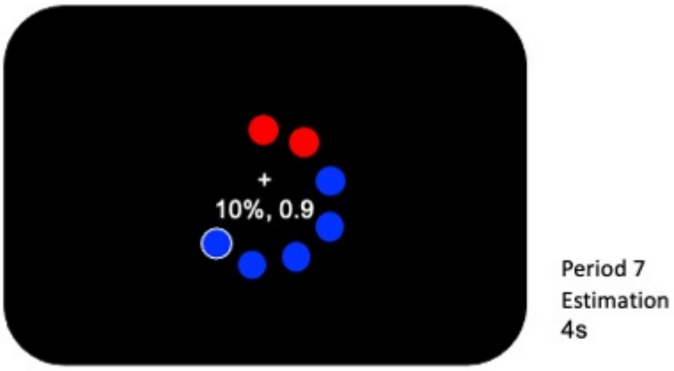
10%, 0.9
Period 7
Estimation
4s

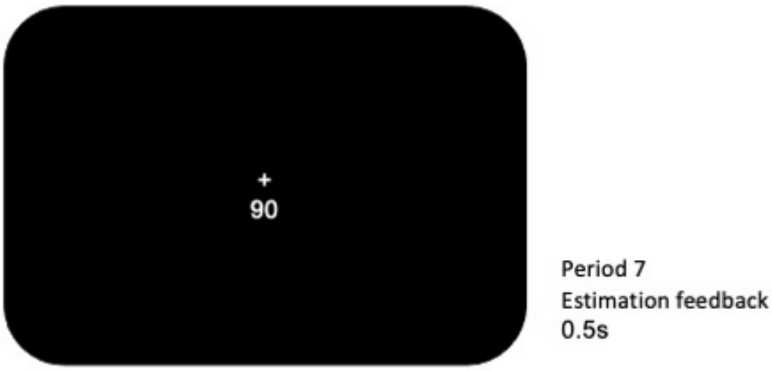
90
Period 7
Estimation feedback
0.5s

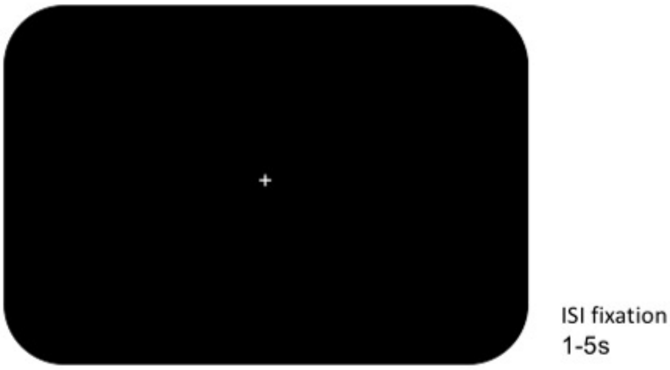
ISI fixation
1-5s

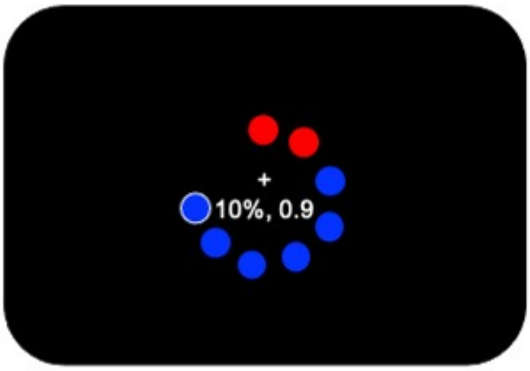


Period 8
Estimation
4s

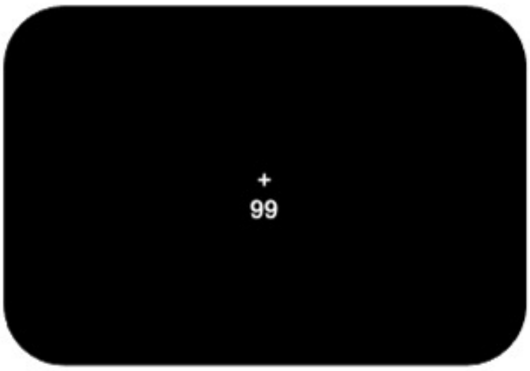


Period 8
Estimation feedback
0.5s

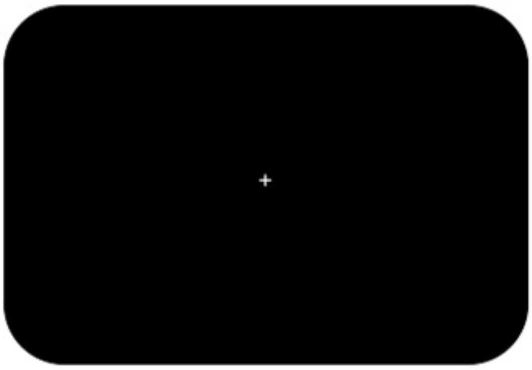


ISI fixation
1-5s

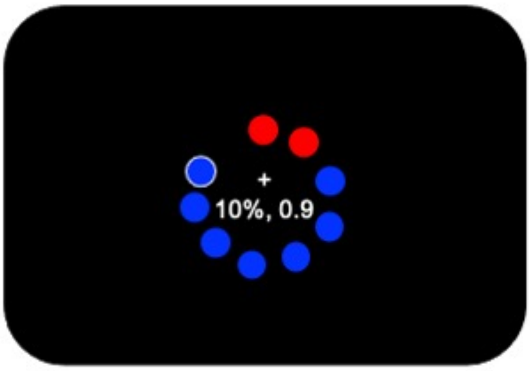


Period 9
Estimation
4s

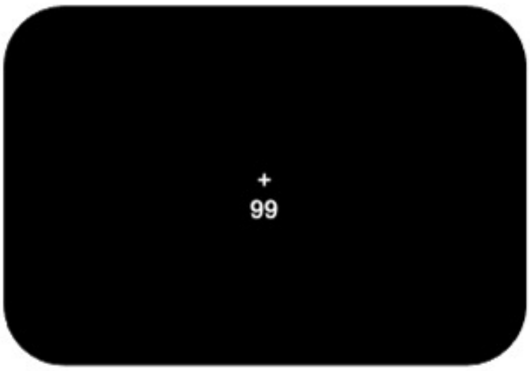


Period 9
Estimation feedback
0.5s

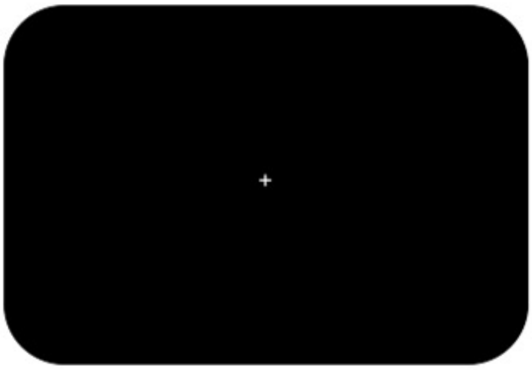

ISI fixation
1-5s

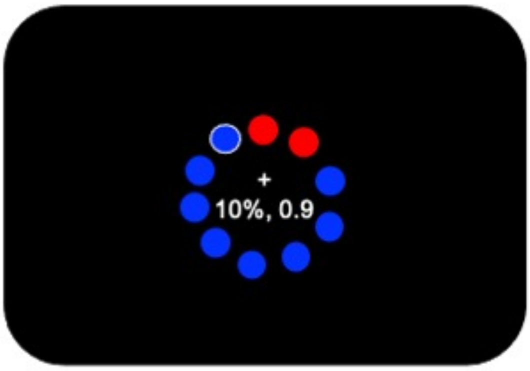


Period 10
Estimation
4s

1

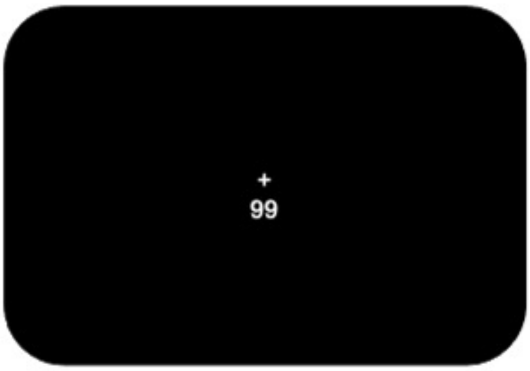


Period 10
Estimation feedback
0.5s

2

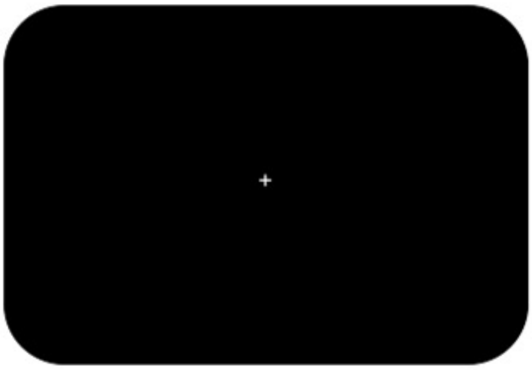

ISI fixation
1-5s

3

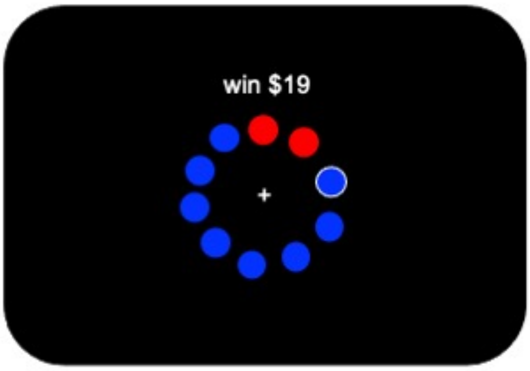


Reward feedback
2s

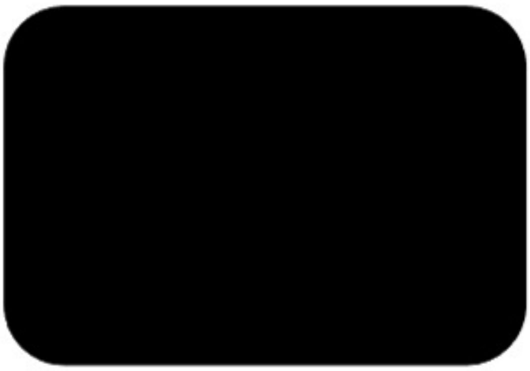

ITI
1-5s
